# Structural basis for preservation of a subset of Topologically Associating Domains in Interphase Chromosomes upon cohesin depletion

**DOI:** 10.1101/2022.10.22.513342

**Authors:** Davin Jeong, Guang Shi, Xin Li, D. Thirumalai

## Abstract

Compartment formation in interphase chromosomes is a result of spatial segregation between eu- and heterochromatin on a few mega base pairs (Mbp) scale. On the sub-Mbp scales, Topologically Associating Domains (TADs) appear as interacting domains along the diagonal in the ensemble averaged Hi-C contact map. Hi-C experiments showed that most of the TADs vanish upon deleting cohesin, while the compartment structure is maintained, and perhaps even enhanced. However, closer inspection of the data reveals that a non-negligible fraction of TADs is preserved (P-TADs) after cohesin loss. Imaging experiments show that, at the single-cell level, TAD-like structures are present *even without cohesin*. To provide a structural basis for these findings, we first used polymer simulations to show that certain TADs with epigenetic switches across their boundaries survive after depletion of loops. More importantly, the three-dimensional structures show that many of the P-TADs have sharp physical boundaries. Informed by the simulations, we analyzed the Hi-C maps (with and without cohesin) in mouse liver and human colorectal carcinoma cell lines, which affirmed that epigenetic switches and physical boundaries (calculated using the predicted 3D structures using the data-driven HIPPS method that uses Hi-C as the input) explain the origin of the P-TADs. Single-cell structures display TAD-like features in the absence of cohesin that are remarkably similar to the findings in imaging experiments. Some P-TADs, with physical boundaries, are relevant to the retention of enhancer-promoter/promoter-promoter interactions. Overall, our study shows that preservation of a subset of TADs upon removing cohesin is a robust phenomenon that is valid across multiple cell lines.

## I. INTRODUCTION

Advances in experimental techniques have provided glimpses of the three-dimensional (3D) organization of chromosomes in diverse species[1–6]. The average (performed over a large number of cells) contact map [4, 6], inferred using chromosome conformation capture technique and related variants (referred to as Hi-C from now on), is a two-dimensional (2D) matrix, whose elements are a measure of the probability that two loci separated by a certain genomic distance are spatially adjacent. The Hi-C experiments on different mammalian cells suggest that there are two major length scales in the organization of interphase chromosomes. On the scale, *L*_C_ ∼ (2-5) Mbp (mega base pairs or Mbp), one observes checkerboard patterns in the contact maps [4], which are thought to be associated with micro-phase separation between the two major epigenetic states, active (A) or euchromatin and inactive (B) or heterochromatin. On the length scale, *L*_TAD_, from tens of kb up to a few Mb, domains, referred to as Topologically Associating Domains (TADs), appear as squares along the diagonal of the contact maps [7, 8]. Contacts are enriched within the TADs, and are suppressed across the boundaries between the TADs. A number of polymer models [9–14] have shown that compartment formations and TADs may be explained using micro-phase separation between A and B type loci. The use of two length scales, *L*_C_ and *L*_TAD_, in characterizing the organization of interphase chromosomes is now entrenched in the field, although there are suggestions that finer sub-TAD structures emerge at kilo base scales [15, 16]. In particular, recent Micro-C experiments have shown that there are fine structures starting from the nucleosome level [17–19], thus establishing the hierarchical organization of interphase chromosomes over a broad range of length scales.

TADs are thought to regulate gene expression by constraining the contacts between target gene and regulatory regions [20, 21]. As a consequence, perturbation or disruption of their integrity such as deletions, duplications or inversions of DNA segments within the TADs, could lead to aberrant gene expression [8, 22–28]. A class of chromatin loops, mediated by the ATP-dependent motor cohesin [29] and the DNA-binding protein CTCF protein (“cohesin-associated CTCF loop”), organizes a subset of the TADs [30]. It is thought that cohesin [29] extrudes DNA loops of varying lengths, which are terminated when the motor encounters the transcriptional insulator CCCTC-binding factor (CTCF) [31]. This implies that cohesin and CTCF are often colocalized at the TAD boundary [4–6, 30, 32, 33].

Several experiments have shown that depletion of the architectural proteins (Nipbl, RAD21, and CTCF) disrupts the organization of interphase chromosomes [26, 34–41].

Schwarzer et al.[34] showed that the removal of the cohesin loading factor, *Nipbl* in the mouse liver cell results in loss of TADs. They concluded that compartment formation, which is independent of cohesin, is a consequence of the underlying epigenetic landscape, while TAD formation requires cohesin. Similarly, it was found that upon removal of cohesin subunit, *RAD21* cohesin-associated CTCF loops and TADs are abolished [26, 39, 40]. Deletion of *RAD21* results in the complete loss of the so-called loop domains [26], which are formed when CTCF colocalizes with cohesin. In contrast, imaging experiments[40] showed that TAD-like structures, with sharp boundaries, at the single-cell level survive even after deleting cohesin. Three headlines emerged from these studies: (i) They reinforce the two-length scale description of genome organization at the ensemble level. (ii) Factors that prevent the association of cohesin with chromosomes globally abolish the TADs and the Hi-C peaks, but preserve (or even enhance) compartmentalization. Experimental studies [4, 26, 34, 39, 40] and polymer simulations [12, 35, 42] have shown that the global epigenetic state determines compartment formation, while the more dynamic TADs, with finite lifetimes[43], require ATP-dependent cohesin. (iii) TAD-like features persist in single cells before and after auxin treatment, albeit with changes in the locations of the sharp domain boundaries.

The results of super resolution experiments [40], at the single-cell level (described above), made us wonder if there is evidence for preservation of TADs at the ensemble level upon cohesin depletion. To this end, we first analyzed the experimental contact maps from mouse liver and HCT-116 cells (human colorectal carcinoma cell line), in the presence and absence of cohesin to assess if TADs are preserved. We discovered that, on an average a fraction of TADs, identified using the TopDom method [44], are retained in chromosomes from both the cell lines (Fig. 1) after removing cohesin. These findings raise the following questions. What is the mechanistic basis for the retention of a small but significant fraction of TADs that are preserved after cohesin loss? Is there a structural explanation for TAD retention at the ensemble level (Hi-C), which would reconcile with the results in super resolution imaging experiments showing TAD-like structures at the single level, even without cohesin?

**Fig. 1.**
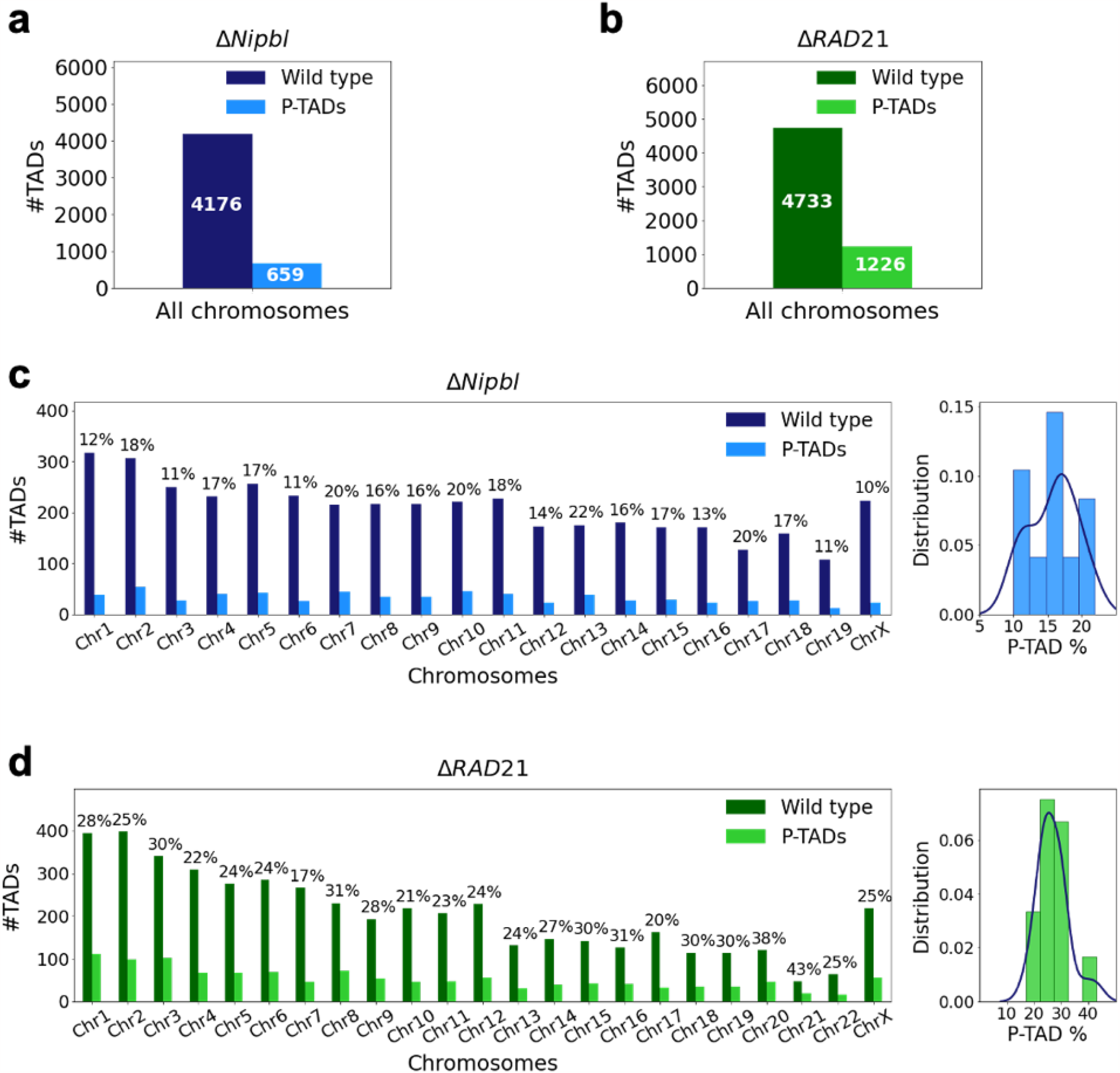
Fate of the TADs in chromosomes upon cohesin deletion. (a) The number of TADs in all the chromosomes, identified by the TopDom method [44], in the wild type (WT) cells and the number of preserved TADs (P-TADs) after deleting cohesin loading factor (Nipbl) in mouse liver. (b) Same as (a) except the experimental data are analyzed for HCT-116 cell before (WT) and after RAD21 deletion. (c) The total number of TADs and the number of P-TADs for each chromosome calculated using the mouse liver Hi-C data. The number above each bar is the percentage of P-TADs in each chromosome. (d) Same as (c) except the results are for chromosomes from the HCT-116 cell line. The percentage of P-TADs is greater in the HCT-116 cell line than in mouse liver for almost all the chromosomes, a feature that is more prominent in the distribution of P-TADs proportions (Right).

We answered the questions posed above by using the following strategy. We first performed polymer simulations for two chromosomes from the GM12878 cell line using the Chromosome Copolymer Model (CCM) [11] with and without loop anchors, which mimics the wild type (WT) and the absence of cohesin-associated CTCF loops, respectively. The major purpose of the CCM polymer simulations is to determine the mechanisms for the emergence of P-TADs. Because the simulations directly generate 3D structures, they can be used to compute average contact maps that can be compared to experiments to determine the accuracy of the CCM. In addition, comparisons of the contact maps with and without cohesin, allowed us to generate the mechanisms for the emergence of P-TADs. Using the polymer simulations of chromosomes from the GM12878 cell line [4], whose organization without cohesin is unknown, we determined that P-TADs arise due to epigenetic switches across TAD boundaries and/or associated with peaks in boundary probabilities, which require knowledge of ensemble of 3D structures.

Informed by the results from the polymer simulations, we analyzed the experimental data from two cell lines [26, 34]. We discovered that epigenetic switch does account for a reasonable fraction (≈ 0.4 in mouse liver and ≈ 0.3 in HCT-116 cell lines) of P-TADs. Rather than perform multiple time-consuming polymer simulations, we generated the 3D structural ensemble using the accurate and data-driven HIPPS method [45] of the chromosomes using the experimental Hi-C data. The analyses using the 3D structures accounted for about 53% of the P-TADs, predicted by theTopDom method [44]. Strikingly, the 3D structures revealed TAD-like structures, at the single-cell level, both in the presence and absence of cohesin, which is in accord with the super resolution imaging data [40]. Our work shows that the effects of cohesin removal on chromatin structures are nuanced, requiring analyses of both the epigenetic landscape as well as 3D structures in order to obtain a comprehensive picture of how distinct factors determine interphase chromosome organization in the nucleus. Our calculations for chromosomes from three cell lines lead to the robust conclusion that a subset of P-TADs is intact after deleting cohesin.

## II. RESULTS

### A non-negligible fraction of TADs are preserved upon removal of cohesin

Experiments [26, 34–37, 39, 40] have shown that deletion of cohesin loaders (*Nipbl in mouse liver, SCC2 in yeast*), cohesin subunit (*RAD21*) abolishes a substantial fraction of both cohesin-associated CTCF loops and the TADs. These observations across different cell lines raise an important question: Do all the TADs completely lose their contact patterns after removal of cohesin? To answer this question, we first analyzed 50kb-resolution contact maps from the two cell lines (mouse liver [34] and HCT-116 [26]) before and after degradation of *Nipbl* and *RAD21*, respectively (see Appendix Analyses of the experimental data for details). Using TopDom [44], we discovered that roughly 659 TADs out of 4176 (16%) are preserved (Fig. 1a) after removing *Nipbl* in the mouse liver cells. In the HCT-116 cells, 1226 TADs out of 4733 (26%)) are preserved (Fig. 1b) upon *RAD21* loss. Fig. 1c and Fig. 1d show that the number of P-TADs depends on the chromosome number. Although the actual number of P-TADs would depend on the TAD-calling protocol, the the finding that a non-negligible fraction is preserved after cohesin depletion is highly significant.

### CCM simulations reproduce wild-type Hi-C maps

To explore the mechanism resulting in P-TADs, we first simulated the chromosome copolymer model [11], CCM (Appendix Fig. 2a). To independently decipher the origins of P-TADs (Fig. 1) in experiments, we calculated the contact maps for Chr13, shown in Appendix Fig. 2 (Chr10 in Appendix Fig. 3,) from the GM12878 cell line. The CCM simulations (Appendix Fig. 2b) reproduce the ubiquitous checker board patterns well. The rectangle in Appendix Fig. 2b represents the border of one such compartment formed primarily by interactions between the B-type loci.

**Fig. 2.**
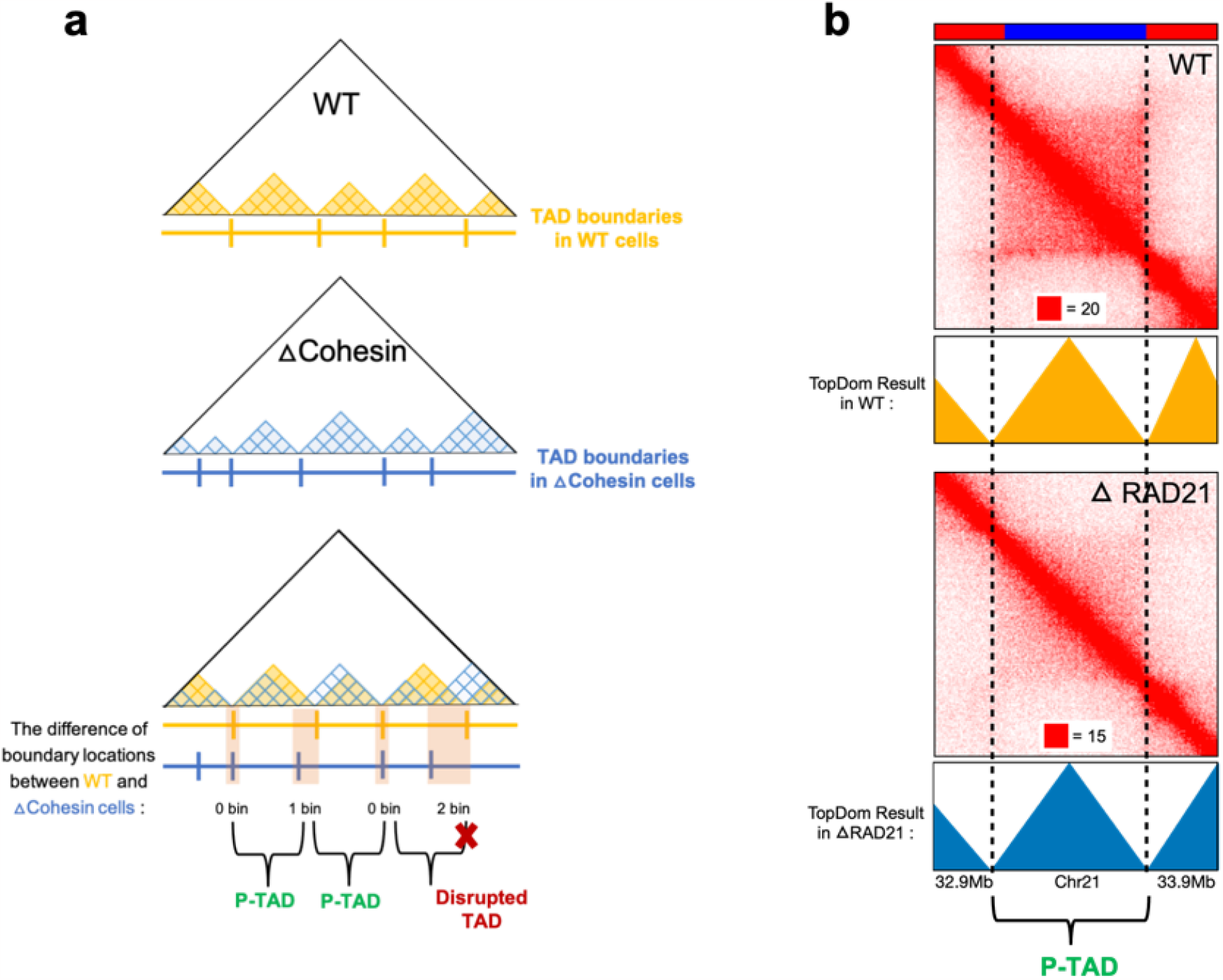
Identification of P-TADs from the contact map using the TopDom method. (a) Schematic representation used to determine the P-TADs. Yellow (Blue) triangles represent the TADs identified using TopDom method in WT (cohesin depleted) contact maps at 50kb resolution.Small square within each triangle represents a single locus (50*kb* size). The boundaries of a TAD detected in the WT contact map within ± one bin (50kb) from a position of boundaries in cohesin depleted cells is deemed to be a P-TAD. (b) P-TAD upon cohesin loss in HCT116 cell. The bar plots above the contact maps show the epigenetic states. Red (Blue) color represents the active (inactive) state, respectively. The TAD between grey dashed lines is preserved upon cohesin loss. The parameter (with red square) displayed at each left bottom indicates the color scale when plotting contact maps used in Juicebox[54].

**Fig. 3.**
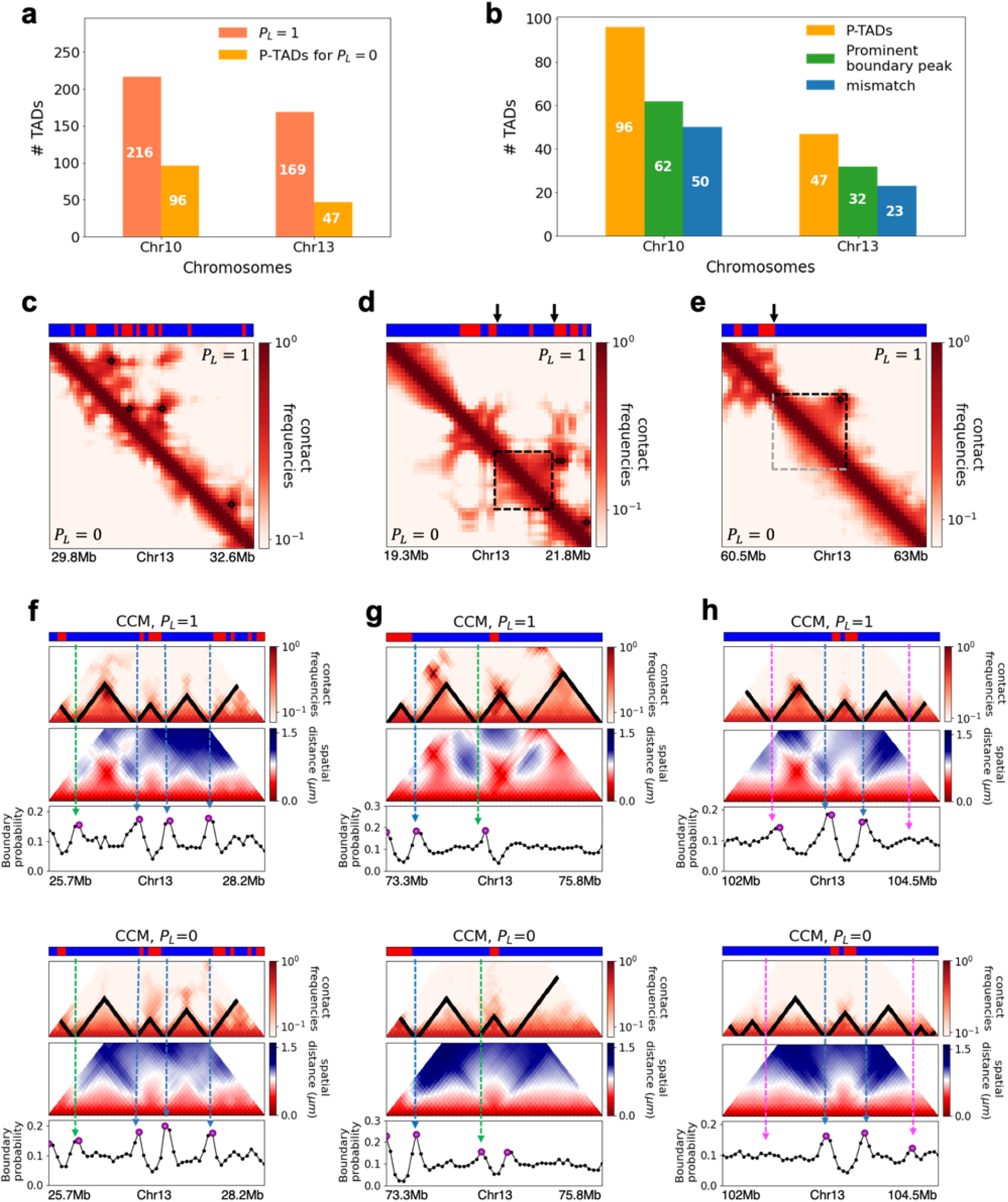
CCM simulations reveal characteristics of P-TADs. (a) The number of TADs in the simulated Chr10 and Chr13 chromosomes for *P*_*L*_=1. The number of P-TADs after CTCF loop depletion (*P*_*L*_=0) is also shown. (b) The number of P-TAD with epigenetic switches (blue) and those identified by the peaks in the boundary probability (green). (c)-(e) Comparison between contact maps for the region of Chr13 with upper (lower) triangle with *P*_*L*_ = 1 (*P*_*L*_ = 0). The black circles at the corner of the TADs are the CTCF loop anchors. The bar above the contact map are the epigenetic states with red (blue) representing A (B) loci. Arrows above the bar show the epigenetic switch. (c) After loop deletion, TAD structures disappear. (d) TAD whose boundaries are marked by epigenetic switches are preserved. (e) TAD lacking at least one epigenetic switch is disrupted after loop loss. (f)-(h) Comparison of the contact map and the mean spatial-distance matrices for the 2.5Mb genomic regions (25.7-28.2*Mbp*,73.3-75.8*Mbp* and 102-104.5*Mbp*, respectively) with (upper) and without (lower) loop anchors. Bottom graph shows the boundary probability, with the high values indicating population averaged TAD boundary. Purple circles in the boundary probability graph represent the preferred boundaries. A subset of P-TADs boundaries match epigenetic switches (blue lines). P-TADs with high boundary probability is in green line. The magenta line describes P-TADs which is not accounted by epigenetic switch or physical boundary in 3D space but are found using the TopDom method.

In order to quantitatively compare the Hi-C data and the simulated contact maps, we transformed the contact maps to Pearson correlation maps, which is used to assess if two loci have correlated interaction profiles (Appendix Fig. 2c). The Kullback-Leibler (KL) divergence between the two probability distributions for the Pearson correlation coefficients, *ρ*_*ij*_s, from simulations and experiments is 0.04 (see Appendix Fig. 2d). We also performed Principal Component Analysis (PCA) on the Pearson correlation matrix to identify the compartment structure. Comparison of the PCA-derived first principal components (PC1) across the Chr13 reveals that A/B compartments observed in the CCM correspond well to those found in the experiments (Appendix Fig. 2e).

We then compared the three-dimensional spatial organization between the simulations and experiments using the Ward Linkage Matrix (WLM), which is based on an agglomerative clustering algorithm. The simulated WLM is calculated from a spatial distance map of the organized chromosome (described in Appendix Fig. 2f-h). We constructed the experimental WLM by converting the Hi-C contact map to a distance map using the approximate relationship [2, 11, 45],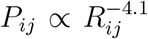. Here, *P*_*ij*_ is the contact probability, and *R*_*ij*_ is the 3D spatial distance between loci i and j (see Appendix Data Analyses). The Pearson correlation coefficient 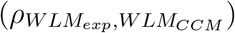 between experimental and simulated WLMs is 0.83 (Appendix Fig. 2h), which establishes the accuracy of the CCM.

Snapshots of TAD structures in Appendix Fig. 2k and n show that they are compact but structurally diverse. The average sizes of the TADs detected using TopDom [44] from Hi-C and simulated contact maps are ∼ 615*kbs* and ∼ 565*kbs*, respectively. Overall the emerging picture of the compartment and TAD structures using different methods are consistent with each other. The results in Appendix Fig. 2 show that the agreement between the CCM simulations and Hi-C data is excellent, especially considering that (i) error estimates in the Hi-C experiments are essentially unknown, and (ii) more importantly, only a single parameter, the inter-loci interaction strength, ϵ, is tuned to fit the experimental contact maps (see Appendix methods sections). Taken together, the results show that the key features of the Hi-C maps for Chr13 (see Appendix Fig. 3 for Chr10 results) are accurately reproduced by the CCM simulations.

### Epigenetic switch accounts for a large fraction of P-TADs

Most of the TADs are not discernible after loop loss, as evidenced by the blurred edges in the contact maps (Fig. 3c). In the CCM simulations of chromosomes from the human GM12878 cell line, a subset of TADs remains even after deleting cohesin-associated CTCF loops (Fig. 3a). The percentages of P-TADs depend on both the resolution of the Hi-C experiments as well as the algorithm used to identify the TADs. By using the same method to analyze both the simulation results and experimental data, it is hoped that the conclusions would be robust.

We used the simulation results to determine the mechanism for the emergence of P-TADs by comparing the results for *P*_*L*_=1 and *P*_*L*_=0. The first observation is that some TADs, even with cohesin-associated CTCF loops, consist mostly of sequences in the same epigenetic state (A or B). Fig. 3d compares the fate of one such TAD in the region (19.3 -21.8 Mb) in Chr13 between *P*_*L*_=1 and *P*_*L*_=0. The highlighted TAD is preserved upon loop loss although the probabilities of contact within this TAD is reduced when *P*_*L*_ = 0 (bottom) compared to *P*_*L*_ = 1 (top). Their boundaries correspond to a switch in the epigenetic state (the sequence location where the change in the epigenetic states occurs, as shown by the two black arrows). In contrast, a TAD in Fig. 3e, that is present in the WT, is abolished when *P*_*L*_=0. The disruption of this particular TAD, lacking in at least one epigenetic switch, occurs because it can interact more frequently with neighboring TADs composed of similar epigenetic states, which in this case is B-type loci. The importance of epigenetic switch in the P-TADs has been noted before [26].

The results in Figs. 3d and e show that a switch in the epigenetic state across a TAD boundary in the WT is likely to result in its preservation after cohesin-associated CTCF loop loss. To test this assertion, we calculated the number of P-TADs that are associated with switches in the epigenetic states (see Appendix Data Analyses and Appendix Fig. 1 for details). We considered the TAD boundary and epigenetic switch as overlapping if they are less than 100kb apart, which is reasonable given that Hi-C resolution adopted here is 50kb. By using 100kb as the cut off, we only consider switch occurrences that exceed two loci. With this criterion, out of 216 (169) TADs calculated using TopDom 50 (23) are P-TADs for Chr10 (Chr13) (vertical blue bars in Fig. 3b in which there are epigenetic switches in the WT).

The P-TADs with epigenetic switches, illustrated in Fig. 3f, shows TADs in the 2.5Mbs region in Chr13. Among the three P-TADs, two of them, whose boundaries are marked by dashed blue lines, have epigenetic switch across the TAD boundary. These two TADs survive after removal of cohesin-associated CTCF loop.

### P-TADs have prominent spatial domain boundaries

Because there are a number of P-TADs that are preserved *even without epigenetic switches* across their boundaries, we wondered if the distance matrix, which requires 3D structures, would offer additional insights about P-TADs upon cohesin-associated CTCF loop loss. Recent imaging experiments[40, 42, 46] revealed that TAD-like domain structures with spatially segregated heterogeneous conformations are present in single cells even without cohesin. The physical boundaries of TAD-like domains, identified from individual 3D structures, vary from cell to cell. They exhibit a preference to reside at specific genomic positions only upon averaging, as found in the Hi-C experiments. The boundary probability at each locus is the proportion of chromosome structures in which the locus is identified as a domain boundary in the 3D space. The locations of prominent peaks in the boundary probability frequently overlap with TADs detected by the population level Hi-C maps.

To explore the relation between P-TADs in ensemble averaged contact maps and preferential boundaries in individual 3D structures of chromosomes, we first calculated individual spatial distance matrices using 10,000 simulated 3D structures which were used to identify the single-cell domain physical boundaries [40]. The physical domain boundaries identified from the 3D structures are the chromosome loci that spatially separate two physical clusters. It is constructed by comparing the spatial distances between a reference locus with the up and down-stream chromosome segments [40]. Specifically, we calculated the median values of pairwise distances between the reference loci and the up-stream loci, and also the median values of pairwise distances between the reference loci and the down-stream. The ratio of these two quantities is defined as boundary strength. If a locus’s boundary strength is above a predefined threshold, this locus is defined as a physical boundary locus. The idea is that a physical boundary has large ratio, as it spatially separates up-stream and down-stream chromatin segments. Based on these boundary positions in individual cells, we define the boundary probability of a locus as the probability (fraction of all individual structures) of this locus being the physical boundary in an ensemble of individual structures. The detailed mathematical definition is provided in Appendix Data Analyses and illustrated in Appendix Fig. 6. We find preferential domains, with high peaks, in the boundary probability along the genomic region as well as variations in single-cell domains both in *P*_*L*_ = 1 and *P*_*L*_ = 0 (see Appendix Fig. 8).

The CCM simulations show that TADs with epigenetic switches across the boundary are likely to be preserved after cohesin-associated CTCF loop loss. Furthermore, Fig. 3f (blue dashed lines) shows that single-cell domain boundaries preferentially reside at the TAD boundaries with epigenetic switches, leading to a prominent signature for the structural ensemble after averaging over a population of cells. Interestingly, the P-TAD has prominent peaks in the boundary probabilities (in both the WT and cohesin-depleted cells), sometimes even without epigenetic switch, at the same genomic position as in the contact map (green lines in Fig. 3f and g). These observations imply that the presence of physical boundaries in the 3D structures may be used to identify P-TADs, especially in instances when there are no epigenetic switches. We should note that the simultaneous presence of peaks in the boundary probabilities in the both WT and cohesin-depleted cells is a signature of P-TADs.

Not all preferential boundaries identified in the distance matrices of the WT cells coincide with the TADs detected using the contact map [40]. There is discordance in the TAD boundaries and high peaks in the boundary probability. The top panel in Fig. 3h (magenta lines) shows that in the (102 - 104.5) Mb range, TopDom predicts that there are three P-TADs after loop loss (see the top panel with P_L_ = 0). There are two prominent peaks in the WT boundary probability whose boundaries coincide with the TADs predicted by TopDom (see bottom panel with *P*_*L*_=1 in Fig. 3h). But the peak height for the third TAD is very small. At best, one can deduce from the boundary probabilities (compare the results in Fig. 3h for *P*_*L*_=1 and *P*_*L*_=0) that the middle TAD is preserved, which would be consistent with the TopDom prediction.

We calculated a standardized Z-score for the boundary probability in the genomic region in order to determine the preferred boundaries in single-cell domains. The number of P-TADs that are accounted for by prominent boundary peaks increases if Z-score is reduced. This implies that some P-TADs detected in the contact maps using TopDom have weak physical boundaries in the 3D structures. We considered the maxima, with Z-score values larger than 0.7, as preferred boundaries in order to determine if P-TADs arise due to the presence of strong physical boundary. With this criterion, we obtained good agreement for the mean length of the TADs detected in the contact map using the TopDom method. The averaged sizes of the TADs in Chr13 using TopDom and boundary probability are ∼565*kbs* and ∼535*kbs*, respectively. Quantitative analysis of the boundary probabilities along the genomic region revealed ≈ 66% of the P-TADs in Chr10 and Chr13 have preferential positioning in single-cell domains (green bars in Fig. 3b). Most P-TADs with epigenetic switches display prominent peaks in the boundary probabilities (≈ 85%).

The primary lessons from the simulations, which form the basis for analyzing the experiments on chromosomes from mouse liver and HCT-116 cell lines, are: (i) switch in the epigenetic state across the TAD boundary is a predominant factor in determining the P-TADs after CTCF loop deletion. (ii) The presence of peaks in the boundary probabilities in both the WT and cohesin-depleted cells, calculated from the the 3D structures, accounts for certain fraction of P-TADs. However, in some instances TopDom predictions (used in (i)) are not compatible with boundaries deduced from 3D structures. (iii) The polymer simulations show that the ensemble of 3D structures provides insights into the consequences, both at the single cell and ensemble averaged level, of depleting cohesin.

### Structural explanation of P-TADs upon cohesin removal from analysis of Hi-C data

In order to assess if the conclusions from simulations, summarized above, explain the experimental data, we first calculated the number of P-TADs whose boundaries have switches in the A/B epigenetic states in mouse liver and HCT-116 cell lines. We assigned chromatin state A (active, red) or B (repressive, blue) by analyzing the combinatorial patterns of histone marks using ChromHMM [47] (see Appendix analyses of the experimental data and Appendix Fig. 7). An average over the 20 chromosomes shows that 280 P-TADs were associated with a switch between A and B epigenetic states upon Δ *Nipbl* in mouse liver (blue bar in Fig. 4a). The corresponding number of P-TADs, averaged over 23 chromosomes, with epigenetic switches is 396 after deleting *RAD21* in HCT-116 (blue bar in Fig. 4b). Not unexpectedly, TADs with epigenetic switches across their boundaries are preserved with high probability after cohesin deletion. We also find that a large number of P-TADs are accounted (green bars in Fig. 4a and Fig. 4b) for by the presence of peaks in the boundary probabilities in the cohesin-depleted cells, which we discuss in detail below.

**Fig. 4.**
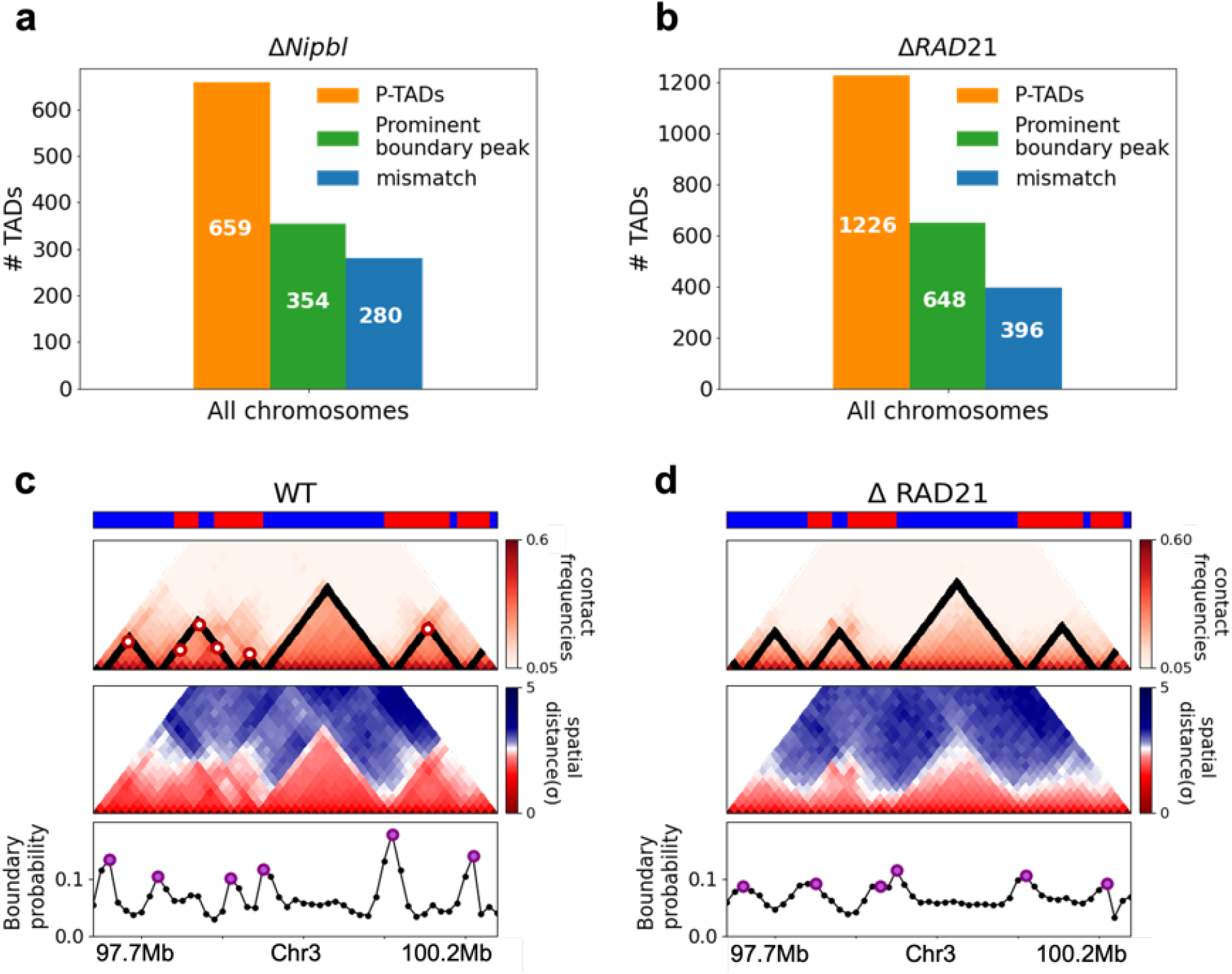
Classification of P-TADs from Hi-C maps from two cell lines and link between boundary probability peak and epigenetic switch. (a) The number of P-TADs in all the chromosomes (orange bar taken from Fig. 1a) that are accounted for by epigenetic switches (blue bar) as well as peaks in the boundary probability (green bar) after *Nipbl* loss in mouse liver. (b) Same as (a) except the analyses is done using experimental data are for HCT-116 cell after *RAD21*. (c) Example of P-TAD in the WT 97.7Mb-100.2Mb region of Chr3 from HCT-116 cell line. The mean distance matrices calculated using the 3D structures is in the middle panel.The dark-red circles at the boundaries of the TADs in the contact maps are loop anchors detected using HiCCUPS [55]. The peaks in the boundary probability (bottom panel) are shown by purple circles. Epigenetic switch coincides with peak in the boundary probability (compare top and bottom panels). Bottom plot shows the probability for each genomic position to be a single-cell domain boundary. (d) Same as (c) except the results correspond to the absence of *RAD21*. Although not as sharp, there is discernible peak in the boundary probability when there is an epigenetic switch after removal of *RAD21*.

We then searched for a structural explanation for P-TADs in the two cell lines (Fig. 4a and Fig. 4b). A plausible hint comes from the the CCM simulations (Fig. 3), which show that boundary probabilities, whose calculations require 3D structures, are good predictors of P-TADs. This implies that peaks in the boundary probabilities should correspond to P-TADs. Similar findings are obtained by analyzing the experimental data (Fig. 4c and d). Additional examples are discussed in Fig. 5. In light of these findings, we wondered if, in general, physical boundaries can be inferred directly using Hi-C data from ensemble experiments, instead of performing tedious polymer simulations. To this end, we used the HIPPS method (see Methods), to calculate an ensemble of 3D structures with the Hi-C contact map as the only input. Several conclusions follow from the results in Fig. 5. (i) Figs. 5a and b show that HIPPS faithfully reproduces the Hi-C contact maps. (ii), Using the ensemble of 3D structures, we calculated the locus-dependent boundary probabilities for both the WT and cohesin-depleted cells. Comparison of the peak positions in the averaged boundary probabilities and the TAD boundaries shows that they often coincide, although there are discordances as well (Fig. 5 and Appendix Figs.12 and 13). (iii) When there is a switch in the epigenetic states, a substantial fraction of P-TADs have high peaks in the boundary probabilities (see Fig. 5 and Appendix Fig. 12). As in the simulations, a large fraction of P-TADs (≈ 67%) have high peaks in the boundary probabilities. Taken together, the results show that the predictions using boundary probabilities and the TopDom method are consistent. (iv) Analyses of the experiments suggest that the epigenetic state as well as the presence of physical boundary in the 3D structures have to be combined in order to determine the origin of P-TADs in cohesin depleted cells.

**Fig. 5.**
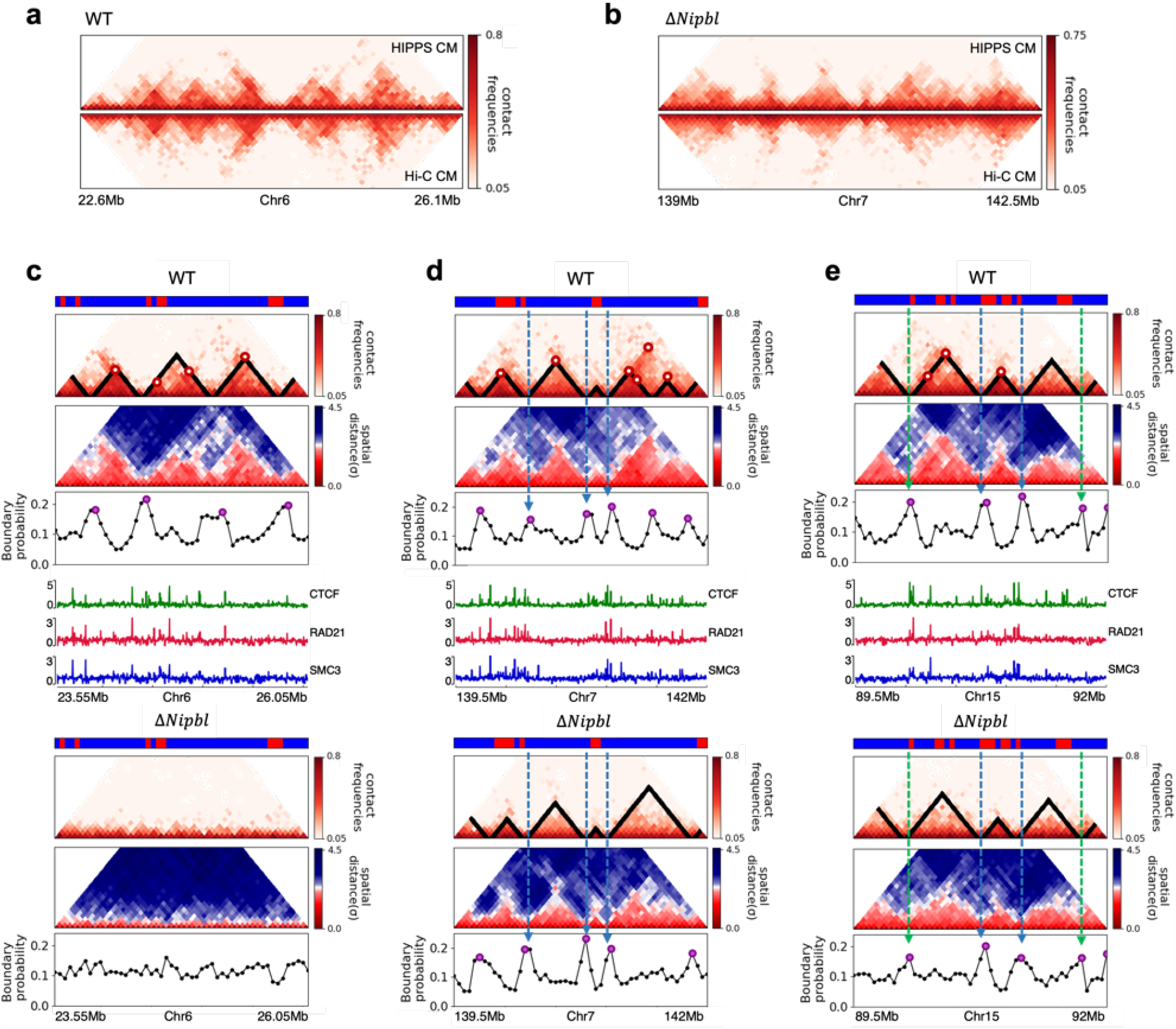
Fate of TADs after Δ*Nipbl* in mouse liver cells. (a)-(b) Comparison between Hi-C (lower) and calculated contact maps (upper) using the 3D structures obtained from HIPPS method for the 3Mb genomic regions (Chr6:22.6Mb-26.1Mb in WT cells and Chr7:139Mb-142.5Mb in *Nipbl* -depleted cells), respectively. The distance threshold for contact is adjusted to achieve the best agreement between HIPPS and experiments. Calculated contact maps are in very good agreement with Hi-C data for both WT and *Nipbl* -depleted cells. (c) Complete Loss (Chr6:23.55Mb-26.05Mb) of TADs in Δ*Nipbl*. (d)-(e) P-TADs (Chr7:139.5Mb-142Mb and Chr15:89.5Mb-92Mb). The plots below the scale on top, identifying the epigenetic states[47], compare 50kb-resolution Hi-C contact maps for the genomic regions of interest with *Nipbl* (upper) and without *Nipbl* (lower). Mean spatial-distance matrices, obtained from the Hi-C contact matrices using the HIPPS method [45], are below the contact maps. The dark-red circles at the boundaries of the TADs in the contact maps are loop anchors detected using HiCCUPS [4]. ChIP-seq tracks for CTCF, RAD21 and SMC3 in the WT cells [34] illustrate the correspondence between the locations of the most detected loop anchors and the ChIP-seq signals. Bottom plots give the probabilities that each genomic position is at a single-cell domain boundary in the specified regions. Purple circles in the boundary probability graph represent the physical boundaries. A subset of physical boundaries in P-TADs coincide with epigenetic switches (blue lines), indicating that the probabilities of contact at these boundaries are small. P-TADs in (e), demarcated by green lines, have high peaks in the boundary probability in the absence of epigenetic switch.

**Fig. 6.**
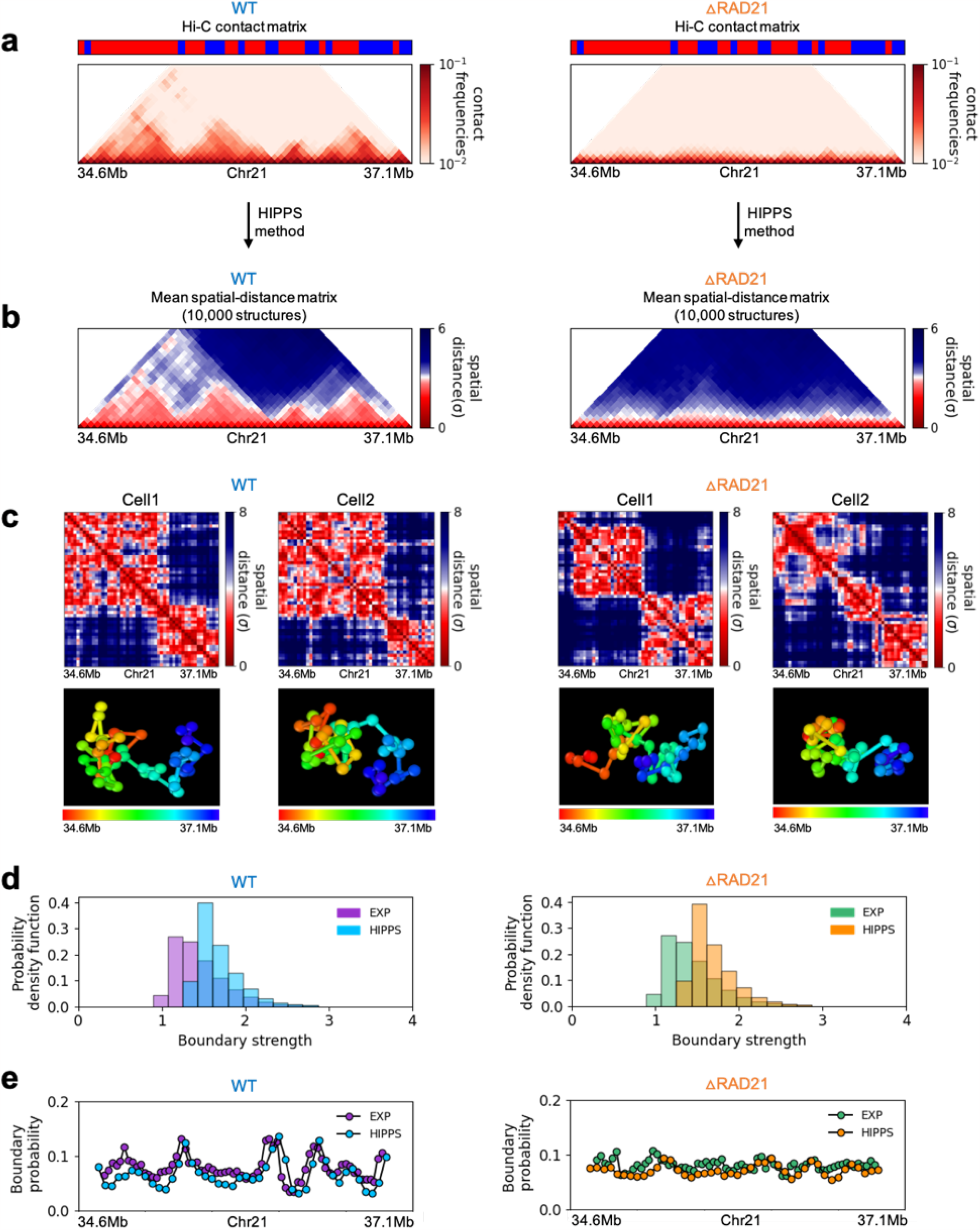

**Fig. 7.**
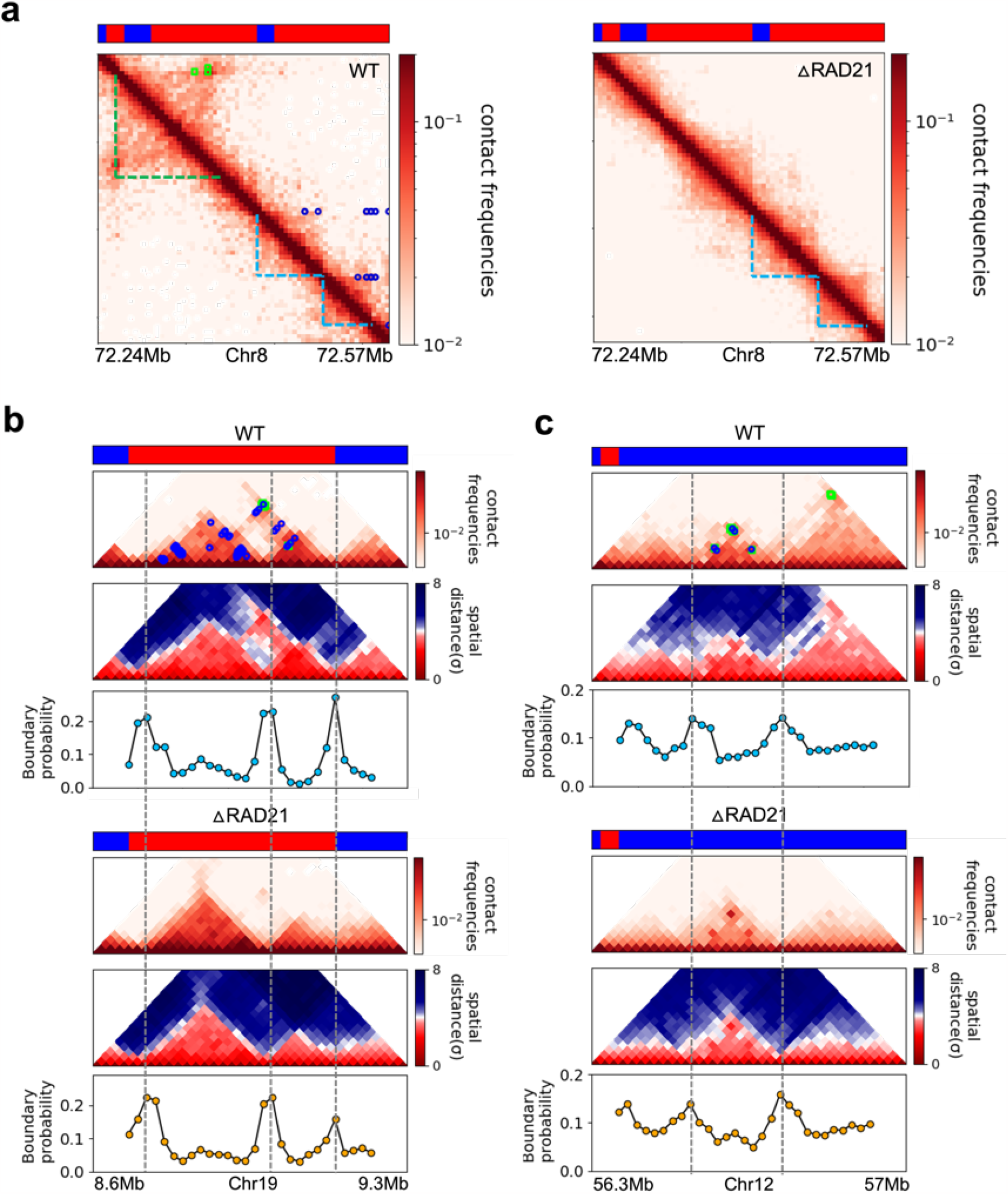
Certain TADs enrichhed in E-P/P-P interactions at the boundary are preserved pon cohesin deletion. (a) Comparison between 5kb Micro-C contact maps in the region (Chr8:72.24Mb-72.57Mb) for the WT (left panel) and cohesin-depleted (right panel) mESC cells[48]. Location of cohesin loops (green square) and EP/P-P (blue circles) plotted in the WT contact maps are from experiments[48]. Bars above the contact map show epigenetic states (Red: Active, Blue: Inactive) annotated based on ChromHMM results[56]. The cohesin-dependent (green dashed lines) and independent (blue dashed lines) TADs are detected in the WT cells using the TopDom method with default parameter (w=5). P-TADs (blue dashed lines) are also found in cohesin deleted cells. (b)-(c) Comparison between 20kb Micro-C contact maps and mean distance maps spanning the regions, Chr19:8.66-9.2Mb and Chr12:56.4-56.9Mb, respectively, in the presence (upper) and absence (lower) cohesin. Bottom graph, below the distance maps, shows the boundary probability calculated from 10,000 3D structures. P-TADs between grey dashed lines are detected using TopDom method (w=5). A P-TAD with high boundary peak, without epigenetic switches, are enriched due to E-P/P-P interactions at the boundaries.

**Fig. 8.**
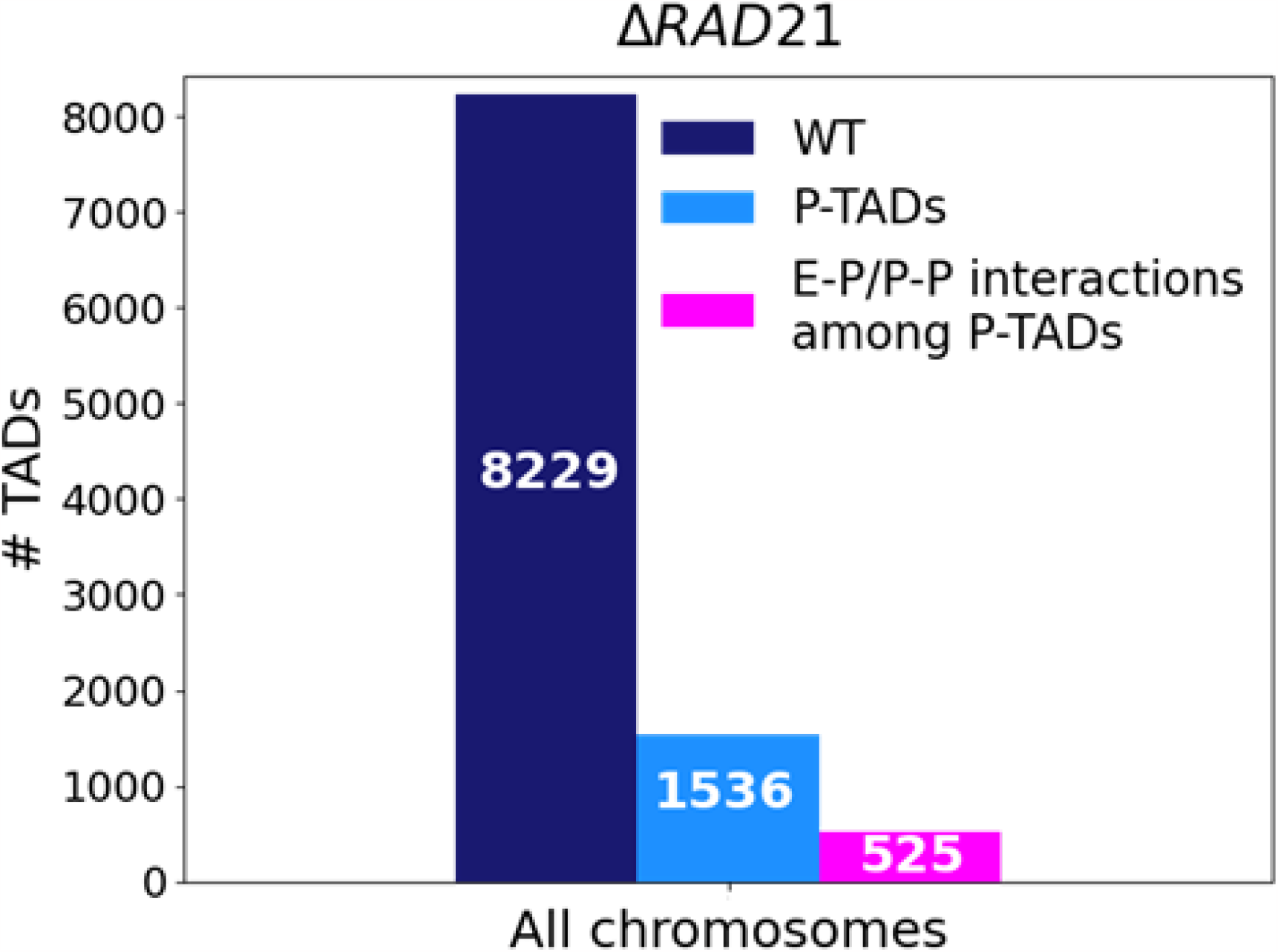
Statistics of the TADs in chromosomes upon cohesin loss using Micro-C contact data. The number of TADs in all the chromosomes in the wild type (WT, dark blue bar), the total number of preserved TADs (P-TADs, light blue bar) after deleting RAD21, and the number of P-TADs whose boundaries coincide withe enhancer-promoter/promoter-promoter (E-P/P-P) interactions (magenta bar) in mESC. About a third of the P-TADs are associated with E-P/P-P interactions.

With the near quantitative agreement with experiments, we performed detailed analyses, based on the epigenetic switches and boundary probabilities for chromosomes from the mouse liver (Fig. 5). The Appendix contains analyses of the experimental data, and the results for HCT-116 cell are given in Appendix Fig. 12. To illustrate different scenarios, we consider the 2.5Mbs regions from Chr6 (Fig. 5c), Chr7 (Fig. 5d), and Chr15 (Fig. 5e). (i) For Chr 6, there are three TADs according to TopDom (Fig. 5c) in the WT. Upon Δ*Nipbl*, these TADs are abolished (compare the top and bottom panels in Fig 5c). The epigenetic track indicates that the region is mostly in the repressive (B) state. Quantification of the boundary probabilities along the 2.5Mb region of Chr6 shows that the TADs also lack physical boundaries upon Δ*Nipbl*. (ii) Examples of P-TADs that satisfy the epigenetic switch criterion are given in Fig. 5d. Using TopDom, we identified several TADs (top panel in Fig. 5d) in this region of Chr7. It is interesting that the boundary probabilities obtained from the HIPPS-generated distance matrices are also large when there is a switch in the epigenetic states. In these examples, both epigenetic switches and boundary probabilities give consistent results (see the dashed blue lines in Fig. 5d). Two TADs in the WT (the ones on the right in the upper panel in Fig. 5d) merge to form a single TAD in the Δ *Nipbl*. This observation is in accord with expectation based on epigenetic switch, whose corollary is that if there is a TAD within a region that contains predominantly A or B type loci they ought to merge upon Δ*Nipbl*. (iii) In the 2.5Mb region of Chr15, there are three TADs in the WT (top panel in Fig. 5e). The first and the third TADs have an epigenetic switch at only one boundary (blue dashed line), and the expectation is that they would not be preserved upon *Nipbl* removal. However, the boundary probabilities show that the TADs have physical boundaries in both, and thus they are preserved. Taken together, the results in Fig. 5 show that by combining the epigenetic switches (Hi-C data is sufficient) and the boundary probabilities (3D structures are required), one can account for a number of P-TADs.

### Single-cell structural change upon cohesin depletion

Finally, we asked whether the HIPPS method captures the 3D structural changes in cohesin-depleted cell at the single-cell level. To this end, we compare the structures obtained using HIPPS with the imaging data [40], which examined the consequences of Δ*RAD21* in HCT-116. We used HIPPS, with Hi-C contact map as input on the same genomic region as in the experiment [40], to generate the 3D structures. The results of our calculations for a 2.5Mbp in Chr21 (34.6-37.1Mb) region from HCT-116 cell line for the WT and ΔRAD21 are presented in Fig. III. The distance maps were calculated from the calculated 3D structures using the HIPPS method (Fig. IIIa). The mean distance maps for the WT and Δ*RAD21* are shown on the left and right panels in Fig. IIIb. Similar results for Chr 4 in mouse liver cell (Chr2 in HCT-116) are displayed in Appendix Fig. 9 (Appendix Fig. 10).

Several conclusions may be drawn from the results in Fig. III. (i) There are large variations in the distance matrices and the domain boundary locations/strengths from cell to cell (Fig. IIIc and d). This finding is in excellent agreement with imaging data [40]. (ii) In both experiments and our calculations, there are TAD-like structures at the single level even after *RAD21* is removed (see the right panel in Fig. IIIc for the theoretical predictions). TAD-like structures in single cells with and without cohesin have also been found using the strings and binders polymer model [12]. (iii) The calculated boundary strength distribution (blue histogram in the left panel in Fig. IIId) for the WT is in reasonable agreement with the measured distribution (purple histogram from [40]). Similarly, the calculated and measured boundary strength distributions for ΔRAD21 cells are also in good agreement (right panel in Fig. IIId). Just as in experiments [40], we find that the distributions of boundary strength are the same in the WT and in cells without *RAD21*. (iv) We also find that the theoretically calculated average locus-dependent boundary probability is in very good agreement with the reported experimental data (compare the curves in the left panel in Fig. IIIe for the WT and the ones on the right for 6.RAD21 cells).

### P-TADs are due to enhancer/promoter interactions

Cohesin is thought to directly or indirectly regulate enhancer-promoter (E-P) interactions. However, a recent Micro-C experiment discovered that E-P and promoter-promoter (P-P) interactions are, to a large extent, insensitive to acute depletion of cohesin [48]. It has been previously shown that E-P/P-P interactions form one or multiple self-associating domains, strips that extend from domain borders and loop-like structures at their intersections at a finer scale [49–51]. Inspired by the recent finding [48], we explored if P-TADs that arise in the absence of epigenetic switches, are required for the maintenance of finer-scale E-P and P-P interactions. We analyzed the Micro-C data [48] in order to shed light on this issue. The left panel in Fig. 7a shows cohesin-associated (green dashed line) and cohesin-independent (blue dashed line) TAD structures (defined using Topdom) in the WT cells. In the latter case, the E-P and P-P loops (blue circles) are at the boundary of the TADs even in the absence of *no epigenetic switch*, implying that it is a domain that is needed for E-P or P-P communication. Interestingly, the TADs were also conserved upon cohesin loss (right panel in Fig. 7a). Analyses of the 3D structures (Fig. 7b) reveal that the TADs with E-P/P-P loops have strong physical boundaries *sans* cohesin. Fig. 7c shows an example of a TAD with both E-P/P-P loops and cohesin/CTCF loops at the boundary in the WT cells that is retained after cohesin deletion, and is associated with prominent boundary peaks. We propose that only a subset of TADs are conserved, potentially for functional reasons.

The statistical analyses of all the P-TADs observed in the Micro-C contact maps across all the chromosomes show that 525 out of 1536 P-TADs have E-P/P-P loops that coincide with their boundaries (Fig. 8). Taken together, our observations suggest that maintenance of E-P/P-P interactions could be the origin of the P-TADs even if there are no epigenetic switches. It is worth emphasizing that these conclusions can only be obtained by analyzing the 3D structures, which we calculated from the Micro-C contact maps using the HIPPS method [45] that does not rely on polymer simulations.

## III. DISCUSSION

By analyzing the experimental Hi-C data [26, 34], we first showed that upon cohesin loss a non-negligible fraction of TADs is preserved, which was not previously noticed. To examine the factors that control the P-TADs, we then performed polymer simulations of two chromosomes in the presence and absence of cohesin-associated CTCF loops from the GM12878 cell line. The polymer simulation results were used to generate hypotheses for the emergence of P-TADs, which were used to explain the major findings reported in the experiments in mouse liver and HCT-116 cells [26, 34]. The simulations showed, switches in the epigenetic states across the TAD boundary account for a large fraction of P-TADs. Even in the absence of epigenetic switches, P-TADs could be preserved, as revealed by the presence of physical boundaries in the 3D structures.

Rather than perform a number of time-consuming polymer simulations, we used the data-driven approach [45] that generates three-dimensional chromosome structures rapidly and accurately using the Hi-C contact maps as input. Analyses of the calculated structures, with and without Nipbl or RAD21, showed that A-type loci form larger spatial clusters after cohesin removal, consistent with enhancement of compartments inferred from Hi-C contact maps (Appendix Fig. 4). Most of the P-TADs, with epigenetic switches in the contact maps, have prominent peaks in the boundary probabilities in both WT and cohesin-depleted cells. An important conclusion from this striking finding is that not only micro-phase separation on the larger scale, *L*_C_ but also some special TADs on a shorter scale, *L*_TAD_ are encoded in the epigenetic sequence.

Remarkably, the conclusion that there are cell-to-cell variations in the distance maps, noted in imaging experiments [40], are affirmed in the calculated 3D structures. This finding is significant because **(1)** only a limited number of loci can be directly imaged whereas Hi-C data can be routinely generated at higher resolution, **(2)** the number of Hi-C data on various cell types and species currently is far greater than that obtained from imaging data.

Let us summarize the novel results, which sets our work apart from previous insightful studies [26, 34, 35]. (1) We showed by analyzing the Hi-C data for mouse liver and HCT-116 cell lines that a non-negligible fraction of TADs is preserved, which set in motion our detailed investigations. (2) Then, using polymer simulations on a different cell type (GM12878), we generated quantitative insights (epigenetic switches as well as structural basis) for the preservation of TADs. Although not emphasized, we showed that deletion of cohesin in the GM12878 cell line also leads to P-TADs, a prediction that suggests that P-TADs may be “universal”. (3) Rather than perform time consuming polymer simulations, we calculated 3D structures directly from Hi-C data for the mouse liver and HCT-116 cell lines, which provided a structural basis for TAD preservation. (4) The 3D structures also showed how TAD-like features appear at the single-cell level, which is in accord with imaging experiments [40]. (5) Finally, we suggest that P-TADs may be linked to the maintenance of enhancer-promoter and promoter-promoter interactions by calculating the 3D structures using the recent Micro-C data [48].

### Comments on the Methods

In order to explore the factors that control the P-TADs, there are two assumptions. (1) The results of the Hi-C experiments are taken at face value in the sense. We view this as an assumption because errors in the Hi-C readouts may be difficult to evaluate even though such experiments are invaluable [52]. (2) The TADs were identified using TopDom, one of many TAD callers. A recent survey [53] shows that, although the finding that TADs are hierarchically organized domains is robust, there are substantial variations in the identification of these domains predicted by different methods. Although TopDom fairs reasonably well in comparison to other methods, there is no guarantee that it identifies the TAD location or the number of TADs accurately. It is only for convenience that we used TopDom as the reference to which the results using the boundary probabilities are compared. (3) Because the prediction of 3D structures using the HIPPS method does not require extensive polymer simulations, it can be used to predict the structural changes for chromosomes that are subject to large-scale perturbations. The excellent agreement between the HIPPS calculations and imaging experiments further bolsters the power of our approach.

## Acknowledgements

We are grateful to Alistair Boettiger for discussions and useful comments. We thank Sucheol Shin, Atreya Dey, and Debayan Chakraborty for useful discussions. This work was supported by the National Science Foundation (CHE 2320256) and the Welch Foundation (F-0019) through the Collie-Welch Regents Chair.

## DATA AVAILABILITY

Data presented in this study is available upon reasonable request to the corresponding author.

## METHODS

We performed polymer simulations for the following reasons: (i) Because all TAD calling schemes are approximate, we evaluated the accuracy of given protocol (TopDom [44] in our study) using the well calibrated CCM. TAD identification in the CCM simulations could be made directly from the 3D structures, thus allowing us to test the validity of the TopDom method. (ii) The combination of 3D structures, assignment of epigenetic states using ChromHMM [47], and accurate calculation of the Hi-C maps using the CCM, were used to determine the origin of P-TADs. (iii) An added bonus is that the polymer simulations on an entirely different cell type (human GM12878) could be used to assess the robustness of the conclusion that certain fraction of TADs are preserved upon deletion of cohesin.

To avoid biases in the formulation of the hypothesis to explain TAD preservation, we simulated chromosomes from the Human Cell line (GM12878), which is different from the cell lines used in experiments [26, 34]. In the main text, we report results for Chr13 (19-115.10 Mbp). The total number of 50*kb* loci is N = 1923, and the total number of loop anchor pairs is 72. To ensure that the results are robust, we also simulated Chr10 (Appendix Fig. 3.

### Chromosome Copolymer Model (CCM) for Chromosomes

We modified the Chromatin Copolymer Model (CCM)[11] in order to simulate full length interphase chromosomes. In the CCM, chromosomes are modeled as a self-avoiding copolymer with A (B) type loci representing the active (repressive) epigenetic state. The connectivity between two nearest neighbor loci (*nn*), *i* and *i* + 1, separated by a distance *r*_*nn*_ = |*r*_*i*_ −*r*_*i*+1_|, is given by a Finitely-Extensible Nonlinear Elastic (FENE) potential,

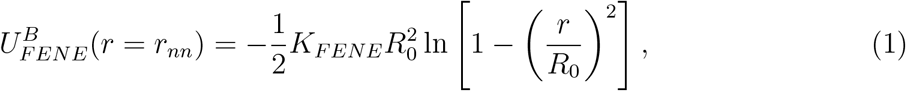

where *K*_*FENE*_ is the spring constant, and *R*_0_ is an estimate of the equilibrium bond distance. Non-bonded interaction between two loci that are not directly connected to each other is given by the Lennard-Jones (LJ) potential,

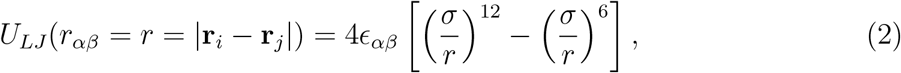

where *α* and *β* can be either A or B. Finally, we used a harmonic potential for the CTCF loop anchors *p* and *q* that are typically stabilized by cohesin. The loop anchor potential is,

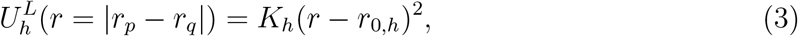

where *K*_*h*_ is the spring constant and *r*_0,*h*_ is the equilibrium length between the CTCF loop anchors. The CCM energy function is,

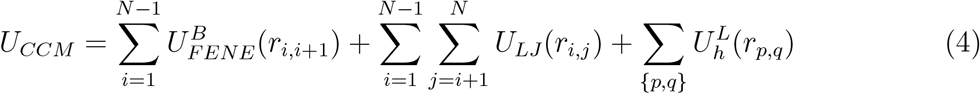

The unit of energy is *k*_*B*_*T*, where *k*_*B*_ is the Boltzmann constant and *T* is the temperature.

We used the CCM simulations in order to deduce the mechanisms for preservation of certain TADs when the loop anchors are deleted. The simulations must reproduce the two major findings [26, 34]: (a) propensity of the A and B loci to segregate should be enhanced upon removal of cohesin; (b) a fraction of TADs should be preserved upon cohesin or cohesin-associated CTCF loop loss. Each locus in the polymer is either A type (active locus is in red) or B type (repressive locus is in blue) (Appendix Fig. 2a). The locus type is determined using the Broad ChromHMM track [57–59]. There are 15 chromatin states, out of which the first eleven are related to gene activity, based on which we group states 1-11 as active state (A), and states 12-15 as repressive state (B). The locations of the CTCF/cohesin-mediated loop anchors, which are fixed in the polymer simulations (cohesin is present), are obtained from the Hi-C data [4] `(GSE63525 GM12878 primary+replicate HiCCUPS with motifs.txt.gz)`. Removal of the loop constraints mimics the absence of cohesin. In the Wild type (WT) simulations, the probability of loop anchor, *P*_L_ being present is unity. To model cohesin depletion, we set *P*_L_ = 0 to assess the impact of deleting the loops on compartments and TADs.

### CCM at 50kb-resolution

In our previous study [11], we used 1,200 bps resolution. Here, we used 50,000 bps (50kb) resolution in order to model the entire length of the chromosomes. To determine the size of each locus, with N_*bp*_ base pairs, we assume[60] that the radius of gyration is 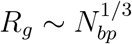. Assuming that a locus, with *σ* _1,200_ and *σ*_50k_, represents a condensed polymer, with 1.2k and 50k base pairs, respectively, we expect that *R*_g,1200_ ∼ (1, 200)^1/3^ and *R*_g,50,000_∼(50, 000)^1/3^. By using this relation, we estimated the size of each locus, _50k_= 3.466 *σ* _1200kb_. Similarly, the mass of the locus at 50kb-resolution is modified as 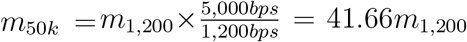, where *m*_1,200_=1. The parameters for the bonding potentials (Eq.1 and Eq.3) at 50kbps resolution of the CCM are given in TABLE 1.

**TABLE 1.**
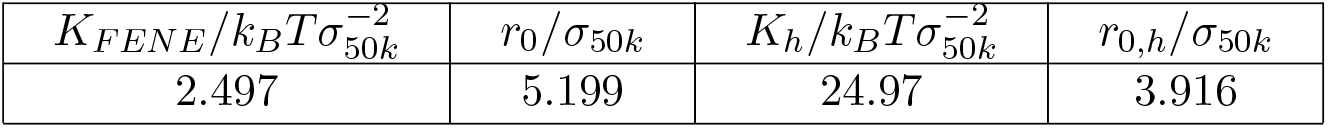
Parameters for bonding potentials.

### Effective energy scales

The creation of CCM was motivated by the experimental observation that active and repressive loci segregate on a few mega base scale. By adopting the Flory-Huggins (FH) theory[60], the spatial segregation between A and B loci is modeled using a weaker A-B attraction compared to A-A and B-B interactions. With the assumption that ϵ_*AA*_ = *ϵ*_*BB*_ *= ϵ*, which is made for simplicity, the only free parameter in the CCM is *ϵ*_*AB*_. By fixing the ratio 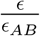 To 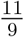, the simulated contact maps are in reasonable agreement with the Hi-C maps. Although a large number of energy functions could reproduce the Hi-C map [10, 13, 61], the CCM is perhaps the simplest copolymer model with only one unknown energy parameter, ϵ.

### Simulations

We performed Langevin Dynamics (LD) simulations using the LAMMPS simulator by integrating the equations of motion,

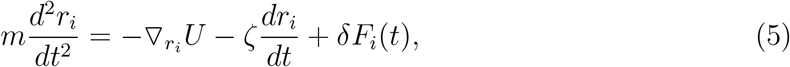

where *r*_*i*_ is the position vector of the *i*^*th*^ locus, and − ∇_*ri*_*U* is the force on the *i*^*th*^ locus, ζ is the friction coefficient that is chosen to be 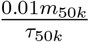. The random force δ *F*_*i*_ (*t*) satisfies ⟨ δ*F*_*i*_(*t*) ⟩ = 0, ⟨ δ *F*_*i*_(*t*) ·*F*_*i*_ (*t*^*′*^) ⟩ = 6ζ*k*_*B*_*T*δ(*t* −*t*^*′*^). We first did simulations using a small time step, 10^6^τ_50k_, with only repulsive pairwise interactions between the loci to avoid numerical instabilities. After a certain number of time steps, the loci associated with the loops are in proximity, and undergo fluctuations around their equilibrium bond distance. At this stage, we increased the time step to 10^2^τ_50*k*_, and turned on the attractive pairwise interactions, and continued the simulations for 10^8^Δ*t*_50*k*_. We then performed LD simulations for an additional 10^8^Δ*t*_50k_ to compute the structural properties. Because we are only interested in equilibrium structures, the values of 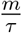 and ζ are irrelevant.

#### Contact map

We calculated the contact frequencies, *C*_*ij*_, between loci i and j by computing the distance, *r*_*ij*_ = |*r*_*j*_ − *r*_*i*_| between them, and counting the number of instances, when *r*_*ij*_ < 1.75 σ _50k_. The set of elements, *C*_*ij*_ constituting the contact map is a 2D representation of the chromosome organization (Appendix Fig. 2(b) and Appendix Fig. 3(a)).

#### Pearson correlation map

To assess the accuracy of the CCM predictions, we calculated the Pearson correlation maps (Appendix Fig. 2(c) and Appendix Fig. 3(b)) by first transforming the simulated contact maps and the Hi-C data to a log_e_ scale. For each element, *C*_*ij*_, we calculated, *Z*_*ij*_, using,

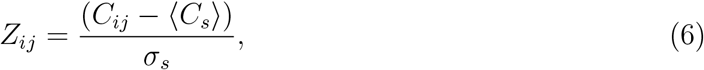

where ⟨ *C*_*s*_⟩ =(1/(*N*−*s*)) ∑_*i*<*j*_ δ(*s*−(*j* −*i*))*C*_*ij*_, and *σ*_*s*_ is the standard deviation associated with *C*_*s*_. The Pearson correlation coefficient (PCC), *ρ*_*ij*_, is calculated between the *i*^*th*^ row, *X*_*i*_, and the *j*^*th*^ column, *Y*_*j*_, associated with the matrix *Z* whose elements are *Z*_*ij*_. The PCC is 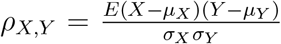 where E denotes expectation, *µ*_*X*_ and *µ*_*Y*_ are the means of *X* and *Y*, respectively, and *σ*_*X*_ (*σ*_*Y*_)
 is the standard deviation of *X* (*Y*).

### Kullback-Leibler(KL) divergence

To measure the difference between two probability distributions that are functions of the same variable x, we calculated the Kullback-Leibler divergence (KL), *D*_*KL*_(*p*(*x*), *q*(*x*)), which is a measure of the information loss when *q*(*x*) is used to approximate *p*(*x*). Here, p(x) and q(x) are the two probability distributions of a discrete random variable x. Using the KL divergence, the difference between the PCC probability distributions obtained from the simulations, *p*^*CCM*^ and experiments, *p*^*EXP*^ were calculated. We define *D*_*KL*_(*p*^*EXP*^, *p*^*CCM*^) as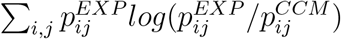 as shown in Appendix Fig. 2(d) and Appendix Fig. 3(c).

#### Ward Linkage Matrix (WLM)

We used the WLM, an agglomerative clustering algorithm method, to reveal the hierarchical organization on different length scales (Appendix Fig. 2((h) and Appendix Fig. 3(f)). In our previous study[11], we showed that the contact probability is inversely proportional to a power of the spatial distance, 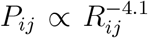. This relationship provides a way to convert Hi-C contact matrix to the spatial distance matrix. We computed WLM with our simulated spatial distance matrix, which is directly calculated in the simulations. To compare with experiments, we converted the Hi-C contact matrix to a spatial distance matrix, **R**_**exp**_ using the relation 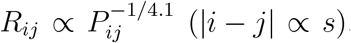. The Ward Linkage Matrix (WLM), **W**, from **R**_**exp**_ and the simulated spatial distance matrix **R**_**sim**_, can be calculated, as described previously [11].

#### Density-based spatial clustering of applications with noise (DBSCAN)

DBSCAN is a clustering algorithm [62] that finds regions of high density by grouping together data points that are in proximity based on spatial distribution. For DBSCAN, two parameters, *Epsilon* and *MinP oints* are required; *Epsilon* is a threshold distance between two loci that is used to classify if they belong to the same cluster whereas *MinP oints* is the minimum number of data points needed to form a dense region. The *MinP oints* can be derived from the dimensions, D in the data points, as *MinPoints D* + 1. We use the recommended value [62] *MinPoints* = 2×D.

The optimal *Epsilon* value is determined using k-distance graph. We set *MinPoints=6* and calculated the distance from every point to the *k*^*th*^ nearest neighbor in each cell. The k-distances are plotted in an ascending order, and a reasonable value corresponds to the maximum curvature (elbow) in this plot. It is likely that optimal values depend on the chromosomes, A/B loci type and the cell type. For example, we found that the optimal *Epsilon* values are 1.7(1.15) σ_*CCM,Chr*13_, 1.0(0.8)σ_*mouseliver,Chr*19_ and 1.6(1.4) σ_*HCT* 116,*Chr*15_ for A (B) loci in Chr13 (CCM), Chr19 (Mouse liver) and Chr15 (HCT116) of both WT and CTCF loops/cohesin depleted cells, respectively (Each represents the average distance between *i* and *i* + 1 loci in the chromosome.). With the optimal parameters, we identified the number of A (B) clusters, N_A_(N_B_) in 10,000 individual structures in the chromosome (see Appendix Fig. 4). In addition, we calculated the size of each cluster, S_A_(S_B_), which is defined as (the number of A (B) loci within the cluster)/(the total number of A(B) loci within the chromosome).

The compartmental strength is enhanced after the removal of CTCF loops (Appendix Fig. 5), indicating that CTCF loop loss leads to an enhanced tendency for micro-phase separation [26, 34–37, 40, 63, 64]. Thus, DBSCAN, a method that relies on 3D structures, and a method that uses only the contact map produces qualitatively consistent picture of strengthening of compartments upon cohesin loss.

#### TAD and P-TAD identification

TopDom [44] is one of many methods used to identify TADs. The average contact frequency around each locus, i between upstream (*i*-*w*+1, *i*-*w*, …, *i*) and downstream (*i*+1, *i*+2, …, *i*+*w*) regions with the free parameter, *w*, is calculated as the value of the binSignal. TAD boundaries correspond to local minima in the binSignal. Subsequently, false detections in the local minima are filtered by using the Wilcox Rank Sum. We used the software package and source codes of TopDom (https://github.com/jasminezhoulab/TopDom) with default parameter, *w*=5. Two aspects concerning the implementation of TopDom should be kept in mind. (i) TopDom results change depending on parameter values. Large w produces big domains, reducing the total number of detected domains. (ii) There are some matrix columns/rows whose contact frequencies sum up to zero. We refer to them as missing bins. We selected only the domains whose boundaries have zero or one missing bin in a 250kb range, since the presence of the missing bin influences contact insulation.

For completeness, let us define preserved TADs (P-TADs). We detected TADs using TopDom [44] based on the Hi-C data. First, *P-TADs* are those that remain in *both* the wild type (WT) cells and cohesin-depleted cells (Fig. 2). If the boundaries between two TADs in cohesin-depleted cells are within *±* 50kb window from the corresponding boundary in the WT, and if there is greater than ≥80% overlap between the WT and cohesin-depleted cells, such a TAD is classified as a P-TAD. Second, *epigenetic switches across TAD boundaries* refer to the alteration of epigenetic state upon going from one TAD to the neighboring TAD (see Appendix Fig. 1). For instance, one TAD consisting of predominantly euchromatin loci with the adjacent TAD comprising largely heterochromatin loci would create epigenetic switches across the boundary. We also used three dimensional structures of chromosomes, with and without cohesin, to calculate boundaries in three dimensions to determine the structural origin of P-TADs.

#### P-TADs with epigenetic switches

The procedure for determining epigenetic switches is schematically shown in Appendix Fig. 1. switches that occupy only one locus (50*kb*) were excluded (see *I* in Appendix Fig. 1). We considered the P-TAD boundary and epigenetic switch as overlapping if they are less than 100kb apart (*II* in Appendix Fig. 1). Finally, P-TADs with epigenetic switches, consisting of <70% of sequences in identical epigenetic states, with epigenetic switches, were filtered out (*III* in Appendix Fig. 1).

#### Boundary strength and boundary probability

To measure the boundary strength [40, 65] for each locus i, we first calculated the median distance values (*L*) of the left three columns, each extending 6-elements below the diagonal, and the median value (*R*) of the right three 6-element columns below the diagonal. Similarly, the median value (*B*) of the right three 6-element columns above the diagonal, and the median value (*T*) of the left three 6-element columns above the diagonal were calculated.

The two boundary strengths, *L*/*R* (start-of-domain boundary strength) and *B*/*T* (end-of domain boundary strength) are computed as defined in Appendix Fig. 6. The local maxima above a defined threshold in the start/end-of domain boundary strengths are identified as te start/end positions of the domain boundary, respectively. This is physically reasonable because at the boundary between two TADs ⟨ p_*ij*_ ⟩ is low which implies that ⟨ r_*ij*_ ⟩ has to be large. Based on the boundary positions in individual cells, we compute the start/end boundary probability for each locus as the fraction of chromosomes in which the corresponding locus is identified as a start/end boundary of a domain. The average of these start and end boundary probabilities for each locus is defined as the boundary probability at the locus (Appendix Fig. 6).

### Appendix A: Analyses of the experimental data

A hypothesis that emerges from the CCM simulations is that the TADs, which are preserved with high probability upon cohesin deletion, have epigenetic switches across the TAD boundary, and are often accompanied by peaks in the boundary probability. In order to test this hypothesis, we analyzed the Hi-C contact map from Schwarzer et al. [34] (mouse liver) and Rao et al. [26] (HCT-116). In each experiment, cohesin loading factor, *Nipbl* and a core component of the cohesin complex, *RAD21* were depleted by employing a liver-specific, tamoxifen-inducible Cre driver and an auxin-inducible degron (AID), respectively. The availability of the wild-type (WT) and cohesin depleted (Δ*Nipbl* or Δ*RAD21*) contact maps allows us to test the hypothesis derived from simulations.

In order to analyze the mouse liver data, we used the wild-type and *Nipbl* -depleted Hi-C contact maps at 50kb-resolution from GEO: GSE93431. The locations of the CTCF loops were determined using the HiCCUPS method in HiCPeaks [4] (https://github.com/XiaoTaoWang/HiCPeaks). HiCCUPS examines each pixel in a Hi-C contact matrix and detects loops by finding the pixels that are enriched relative to local neighborhoods (pixels to its lower-left, pixels to its left and right, pixels above and below, and pixels within a doughnut-shaped region surrounding the pixel of interest). We obtained the locations of the 3,301 loop anchors for 20 chromosomes from the wild type Hi-C contact maps. Epigenetic landscape is determined using the Broad ChromHMM track for mouse liver [66] (https://github.com/gireeshkbogu/chromatin_states_chromHMM_mm9). Among the 15 chromatin states in the track, we assigned states 1-10 and 15 to be in the active state (A) because they are related to gene transcription. States 11-14 correspond to heterochromatin, and hence are taken to the repressive (B).

For the HCT-116 cell, we used 50kb-resolution Hi-C map for untreated and *RAD21* -depleted cells obtained from GEO: GSE104334. Chromatin state characterization was performed using ChromHMM[47] considering twelve histone modifications ChIP-seq data (H3K27ac, H3K9ac, H3K27me3, H3K36me3, H3K79me2, H3K9me2, H3K9me3, H4K20me1, H3K4me1, H3K4me2, H3K4me3, and CTCF) that are available in the ENCODE Project Consortium. Chromatin functional states are annotated based on the study by Moudgil et. al. [67]. For this cell line, we assigned states 1-9 and 15 to be in the the A (active) because they are related to gene transcription. States 10-14 are repressed, and hence may be classified as repressive (B) (Appendix Fig. 7). Locations of the chromatin loops for HCT 116 from Hi-C contact maps were determined using HiCCUPS in juicer [55] (https://github.com/aidenlab/juicer) following the procedure from Rao at al [26]. Using this procedure, we detected 3,624 loop anchors for 23 chromosomes from wild type Hi-C contact maps.

#### Calculation of the 3D structures of chromosome from Hi-C contact maps

In order to identify the A/B clusters and calculate the boundary probability, we need the 3D coordinates of the loci. We used the HIPPS (Hi-C-polymer-physics-structures) [68] method, which uses the Hi-C contact map as the input, to generate an ensemble of three-dimensional chromosome structures. In the HIPPS method, the mean distance matrix is generated using polymer physics using a power-law relation between the mean contact probability, ⟨ *p*_*ij*_ ⟩ between loci *i* and *j* and the average spatial distance, ⟨*r*_*ij*_⟩. An ensemble of 3D structures is calculated by using the mean distance matrix as a constraint using the principle of maximum entropy. We generated an ensemble of 10,000 individual 3D structures by applying the HIPPS method to Hi-C contact map for each chromosome for the two cell lines. The 3D structures are used to calculate the boundary positions (see below) to identify potential single-cell domains, which complements the TopDom analysis.

### Appendix B: Effect of *RAD21* removal

Single-cell studies [12, 40, 42] on a 2.5Mb region (Chr21:34.6-37.1Mb) in HCT116 cell showed that several pronounced TAD structures detected in the WT cells were eliminated if *RAD21* is degraded. This could be predicted by the EMH because the degraded TADs are mostly composed of active (A) loci. Our analysis for the same region also shows flat domain boundary probability (Appendix Fig. 12a). In contrast, Appendix Fig. 12b reveals that preferential TAD boundaries persist, and are retained despite *RAD21* loss, which is associated with epigenetic switches. Furthermore, some TADs without epigenetic switches are preserved, and could be identified based on the presence of physical boundaries in the 3D structures (green lines in Appendix Fig. 12c).

### Appendix C: Corner dots in P-TADs

Our analyses of the experimental data show that TADs whose borders correspond to both epigenetic switch and CTCF loop anchors in the WT cells are preserved after removal of cohesin. However, not all P-TADs have loop anchors at their boundaries (corner dots) in the WT cells (see Appendix Fig. 11). Out of the 280 (396) P-TADs with epigenetic switches in mouse liver (HCT-116) chromosomes, 117 (170) did not have corner dots at their boundaries before deleting Nipbl (RAD21). This suggests that some TAD boundaries are formed in the absence of corner dots. It is the underlying epigenetic landscape that is the predominant factor in the formation of such domain boundaries.

### Appendix D: Loss of cohesin-associated CTCF loops leads to an increase in the degree of compartmentalization

The first experimental finding is that cohesin knockdown results in preservation or even enhancement of the compartment structure[26, 34, 36, 39, 40]. To ensure that CCM reproduces this finding, we first explored how cohesin-associated CTCF loop deletion affects compartmentalization by performing simulations without loop anchors, which is implemented by setting *P*_*L*_ = 0 (see Methods). We find that the plaid patterns persist after the deletion of the loops (Appendix Fig. 4a). Visual comparison between the two contact maps in Appendix Fig. 4a shows that this is indeed the case. Moreover, the rectangles in Appendix Fig. 4a also show that finer features, absent when *P*_*L*_ is unity, emerge when *P*_*L*_=0, which is an indication of increase in the compartment strength. Pearson correlation maps (lower panel in Appendix Fig. 4a) illustrate the reduction in the contact profiles between A and B loci (dark-blue color) after loop loss. There is an enhancement in interactions between same type loci (A-A or B-B) (dark-red color). The results in Appendix Fig. 4a confirm the experimental finding that disruption of the loops not only creates finer features in the compartment structures but also results in increased number of contacts between loci of the same type, which is simultaneously accompanied by a decrease in the number of A-B interactions.

We then investigated if the enhancement of compartments observed in the contact maps has a structural basis. An imaging study[69] showed that active loci form larger spatial clusters in cohesin depleted cells. To probe whether this finding is reproduced in our CCM simulations, we first calculated the spatial distance matrix using 10,000 simulated 3D structures. For individual matrices, with and without cohesin-associated CTCF loops, we identified A (B)-dense regions as A (B) clusters, respectively, using the DBSCAN (Densitybased spatial clustering of applications with noise) method [62]. Appendix Fig. 4b shows the number of clusters obtained using DBSCAN. The number of A clusters varied between individual structures, which reflects the heterogeneity in the chromosome organization. On an average, there are about five A clusters for *P*_*L*_ = 1, which decreases to four with *P*_*L*_ = 0, which is one indication of the increase in the compartment strength. We also calculated the size of the clusters, which is the average fraction of loci in each cluster in individual structures (right panel in Appendix Fig. 4c). On an average, the size of A clusters is greater after loop loss. This implies that active clusters merge in space to form larger and more connected clusters upon deleting the loops. In contrast, there is no change in the number and size of B clusters (see Appendix Fig. 4a). Taken together, our observations for A and B clusters show that the A loci form larger clusters upon loop loss, which leads to stronger segregation between A and B loci after the loops are deleted.

In the Appendix Fig. 4, we used the DBSCAN method to demonstrate the enhanced compartmentalization upon cohesin deletion. In order to assess if our method produces results that are consistent with other techniques, we also calculated the compartment strength used in previous studies [70, 71]. In this method, chromosomal interaction frequencies are first normalized by the average interaction frequency at a given genomic distance, s. Then the distance corrected interaction frequencies are sorted based on the PC1 value for each locus *i*, which we calculated from the contact map. Finally, the frequencies were aggregated into 50 bins to obtain the saddle plot (Appendix Fig. 5) in which the top left corner (B-B) indicates the frequency of interaction between the B compartments, while bottom right (A-A) represents the contact frequency between the A compartments. Top Right and Bottom Left corners are interaction frequencies between A and B compartments. The strength of the compartment is calculated using, ((AA) + (BB)) / ((AB) + (BA)). The values used for this ratio were determined by calculating the mean value of 20% bins in each corner of the saddle plot. We used cooltools for saddle plot calculations (https://github.com/open2c/cooltools).

#### Relation to experiments regarding the compartmentalization

Besides being consistent with imaging experiments[69], we wondered whether the structural changes inferred from the Hi-C maps from the mouse liver and HCT-116 cell lines liver [26, 34] could be used to demonstrate the strengthening of compartments upon deleting cohesin. We calculated an ensemble of 10,000 3D structures for chromosomes from the two cell lines using the HIPPS method [45]. Consistent with the simulation results, we also observed variations in the spatial organization of A and B loci in individual structures in WT and cohesin depleted cells. DBSCAN analysis identified a smaller number A/B clusters with larger size in ΔNipbl (ΔRAD21) cells compared to WT cells, as shown in Appendix Fig. 4d for Chr11 and Chr19 (see Appendix Fig. 4e for HCT-116 results).

To provide additional insights into the calculated conformations, we constructed Pearson correlation matrices using contact maps calculated from the 3D structures. Appendix Figs.4f and g show that cohesin loss induces stronger plaid patterns, which is consistent with simulation results (lower panel of Appendix Fig. 4a) for Chr13 from GM12878 cell line. Comparison of the 3D structural changes with and without cohesin shows that the micro-phase separation between active and inactive loci is strengthened, with larger A and B physical clusters upon depletion of cohesin, which accords well with Hi-C experiments [69].

**Appendix Fig. 1.**
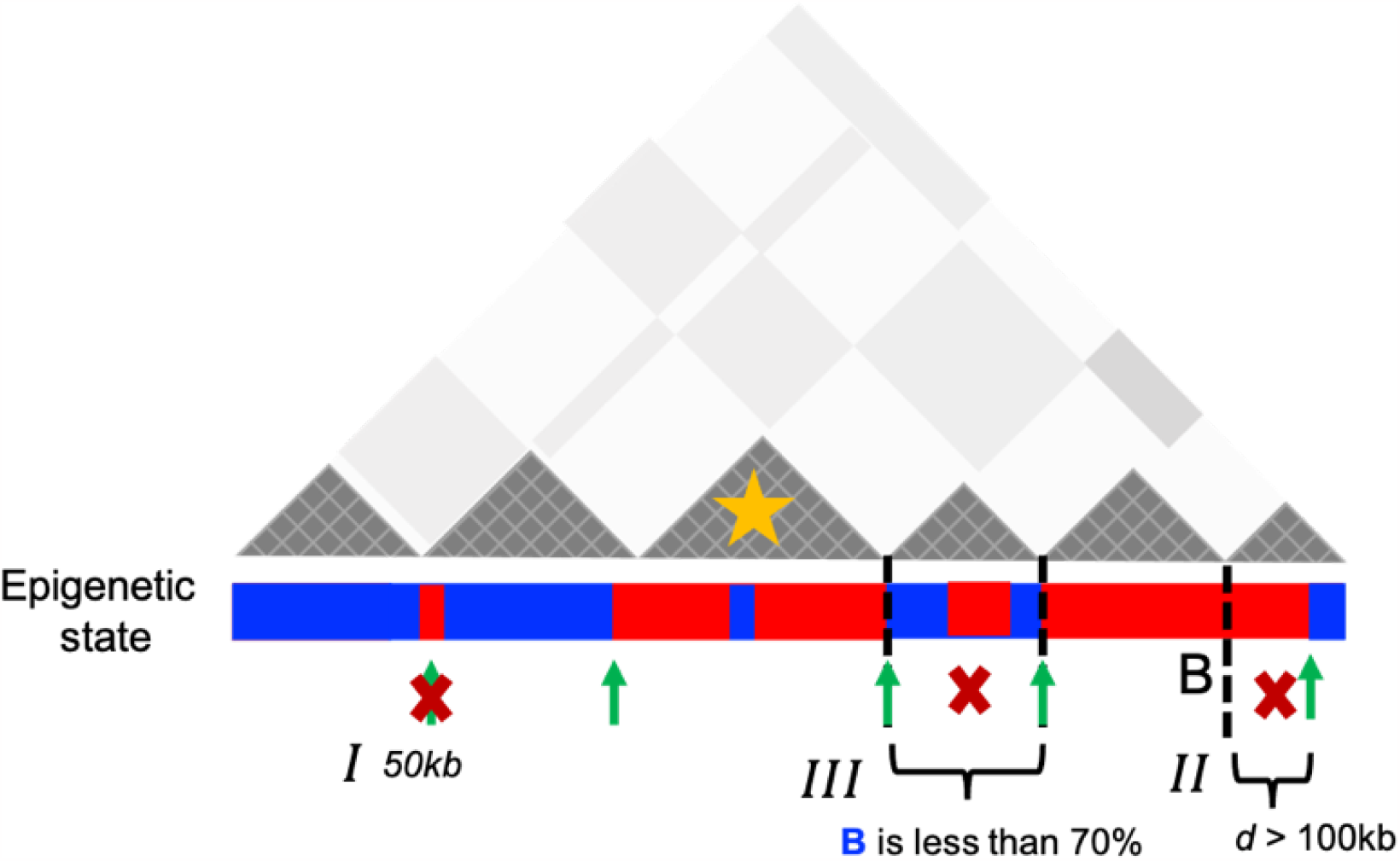
Schematic representation used to identify the P-TADs with epigenetic switches: Dark grey triangles represent the P-TADs in contact map. Small square within each triangle represents a single locus (50*kb*). Red (Blue) color indicates the active (inactive) state in the bar below the contact map. A transition between A and B epigenetic states is referred as to epigenetic switch (Green arrows). We examined if each P-TAD has an epigenetic switch at the boundaries *±* 100kb (*II*). If P-TADs have only one locus (50*kb*) switch near their boundaries (*I*) or comprise *<* 70% of sequences in identical epigenetic state (*III*), they are excluded. The TAD (yellow star) is a P-TAD with epigenetic switch at the TAD boundary.

**Appendix Fig. 2.**
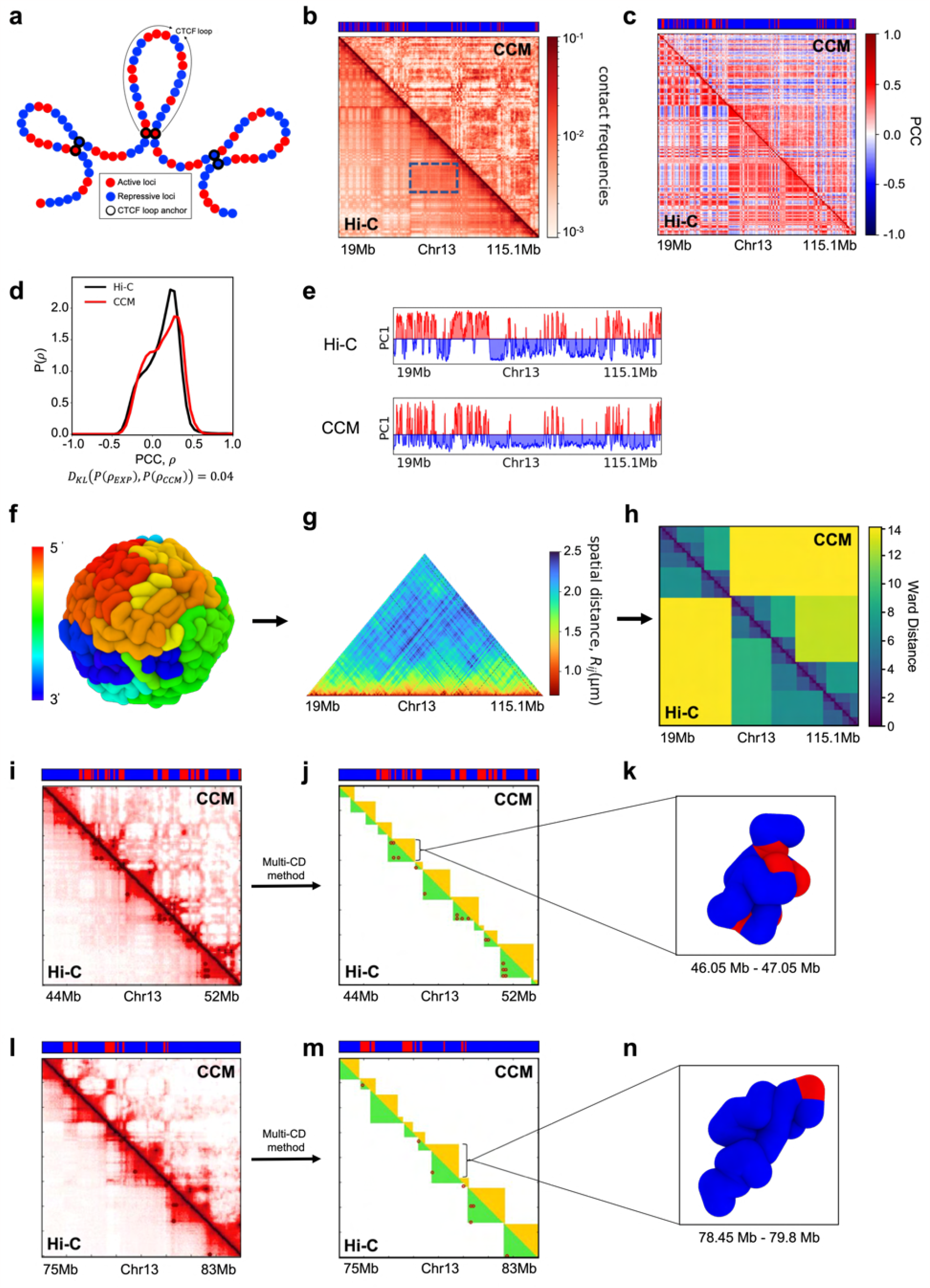
CCM simulations for chromosome 13 (Chr13) from the GM12878 cell line: (a) In the CCM, red (blue) spheres represent active (repressive) loci. The black open circles are the CTCF loop anchor locations. (b) Comparison of the simulated (*P*_*L*_ = 1, top half)) and Hi-C contact maps (bottom half). The bar above marks the epigenetic states with red (blue) representing active (repressive) loci. The values of the contact frequencies, converted to a *log* scale, are shown on the right. (c) Comparison between the Pearson correlation maps consisting of *ρ*_*ij*_ for all loci pairs from simulations (top half) and experimental data (bottom half). The scale for the Pearson Correlation Coefficient (PCC) is on the right. (d) Distribution of the PCC, *ρ*_*ij*_ for all (*i, j*) pairs from simulations and experiment (1 is positive correlation, 0 is no correlation, and -1 corresponds to anti-correlation). The Kullback-Leibler, *D*_*KL*_, value between CCM prediction and experiment is small. (e) First eigenvector values (PC1) from Principal Component Analysis (PCA) using the correlation matrix for CCM. The compartment A and B are defined by positive (red) and negative (blue) values. (f) Snapshot of the folded Chr13. The color corresponds to genomic distance from one end point, ranging from red to green to blue. (g) Ensemble averaged distance map obtained from simulations. (h) Ward Linkage Matrix (WLM) comparison between simulations and the one computed using Hi-C data. The PCC between the two distance matrices is ∼ 0.83, indicating reasonable agreement between simulations and experiments. (i) Contact map for the 8 Mbp region ((44-52)Mb) with the upper (lower) triangle corresponding to simulations (experiments). (j) On the right is an Illustration of the TADs, identified using the Multi-CD method [72]. The dark-red circles are the positions of the loop anchors detected in the Hi-C experiment, which are formed by two CTCF motifs. A subset of TADs is defined by the CTCF loops, whereas others are not associated with loops. These could arise from segregation between the chromatin states of the neighboring domains in certain experimental studies [26, 73, 74]. The average sizes of the TADs detected using Multi-CD method from Hi-C and simulated contact maps are ∼750*kbs* and ∼700*kbs*, respectively. (k) Snapshot of the TAD, marked in (j). (m) Same as (j) except the TADs were calculated for the region ((75-83)Mb) in (l).

**Appendix Fig. 3.**
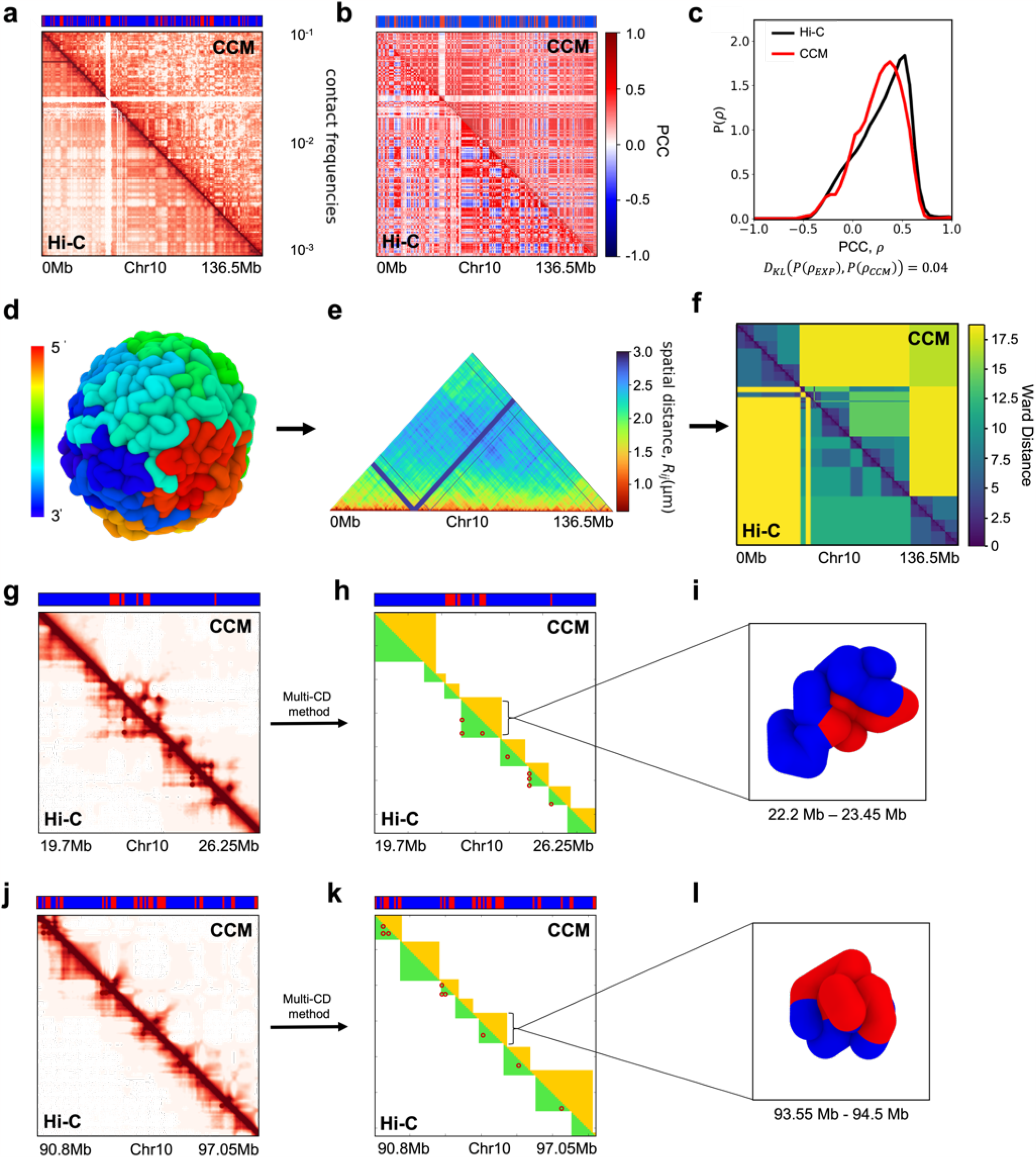
Organizational features of Chr10 from human cell line GM12878: (a) Comparison between the simulated contact map (*P*_*L*_ = 1.0, top half) and Hi-C experiments (bottom half). The bar above the contact map shows the epigenetic states with red (blue) representing active (repressive) loci. (b) Experimental (lower triangle) and the simulated (upper triangle) Pearson correlation maps. (c) The distribution of the PCC, *ρ*_*ij*_ for each pair of (*i, j*) from simulations and experiment. The value of the KL divergence at the bottom is obtained by comparing the distributions obtained in the simulations and experiments. (d) A conformation of the folded Chr10 (N=2,712) obtained using the CCM simulations. The colors correspond to genomic distance from the 5^*0*^ to 3^*0*^ end. (e) Ensemble averaged distance map calculated using the simulated structures. (f) Experimental (lower triangle) and the simulated (upper triangle) WLMs. The PCC between the two WLMs is ∼ 0.75. The agreement between simulations and experiments is fair. (g) Hi-C map for the region (19.7-26.25) Mb with the upper (lower) triangle corresponding to simulations (experiments). (h) Right is an illustration of the TADs. The dark-red circles are the positions of the loop anchors detected in the Hi-C experiment, formed by two CTCF motifs. (i) Snapshot of the TAD, marked by the black line in (h). (k) Same as (h) except the TADs were calculated for a region (90.8-97.05)Mb in (j). The diversity of TAD structures is apparent.

**Appendix Fig. 4.**
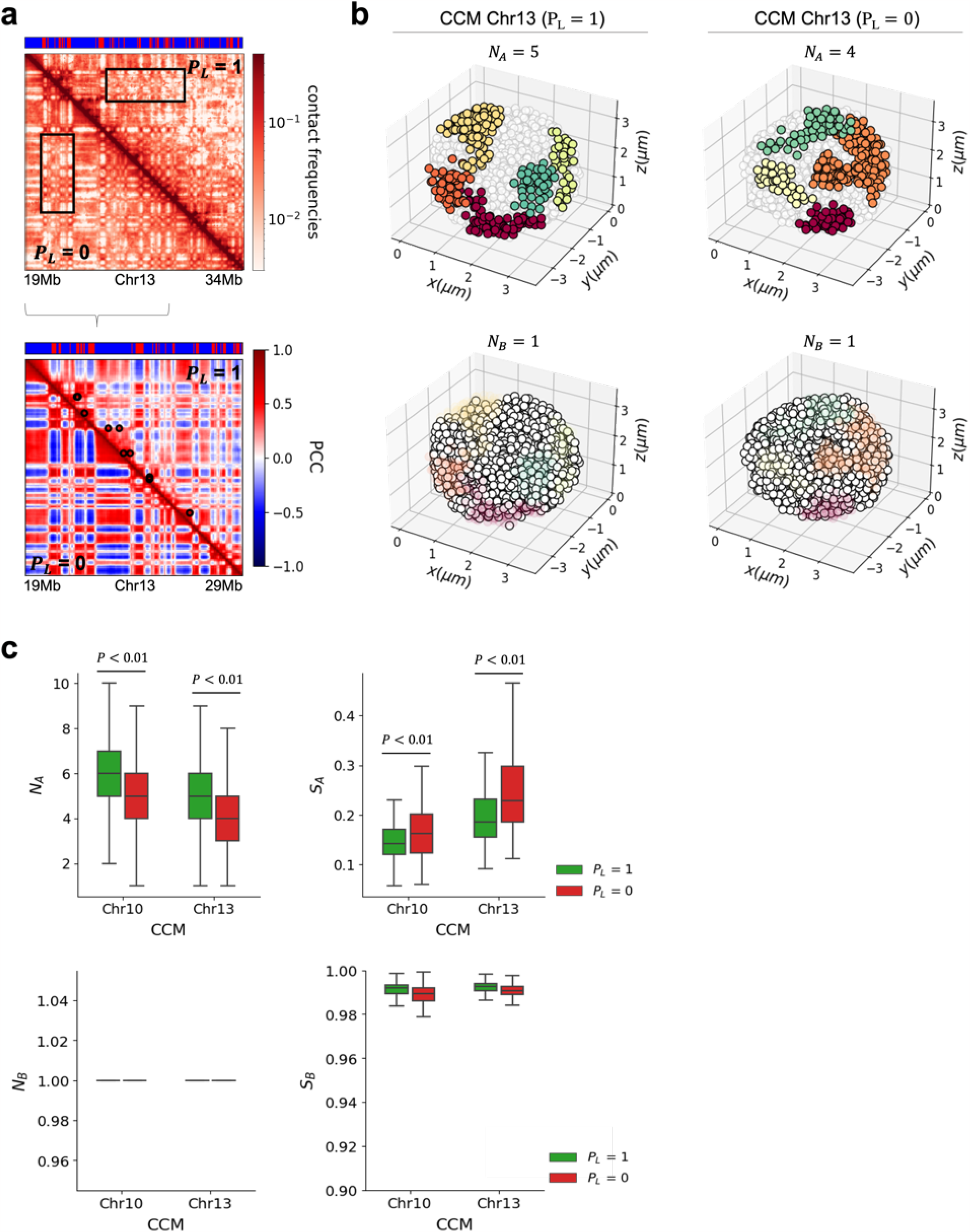
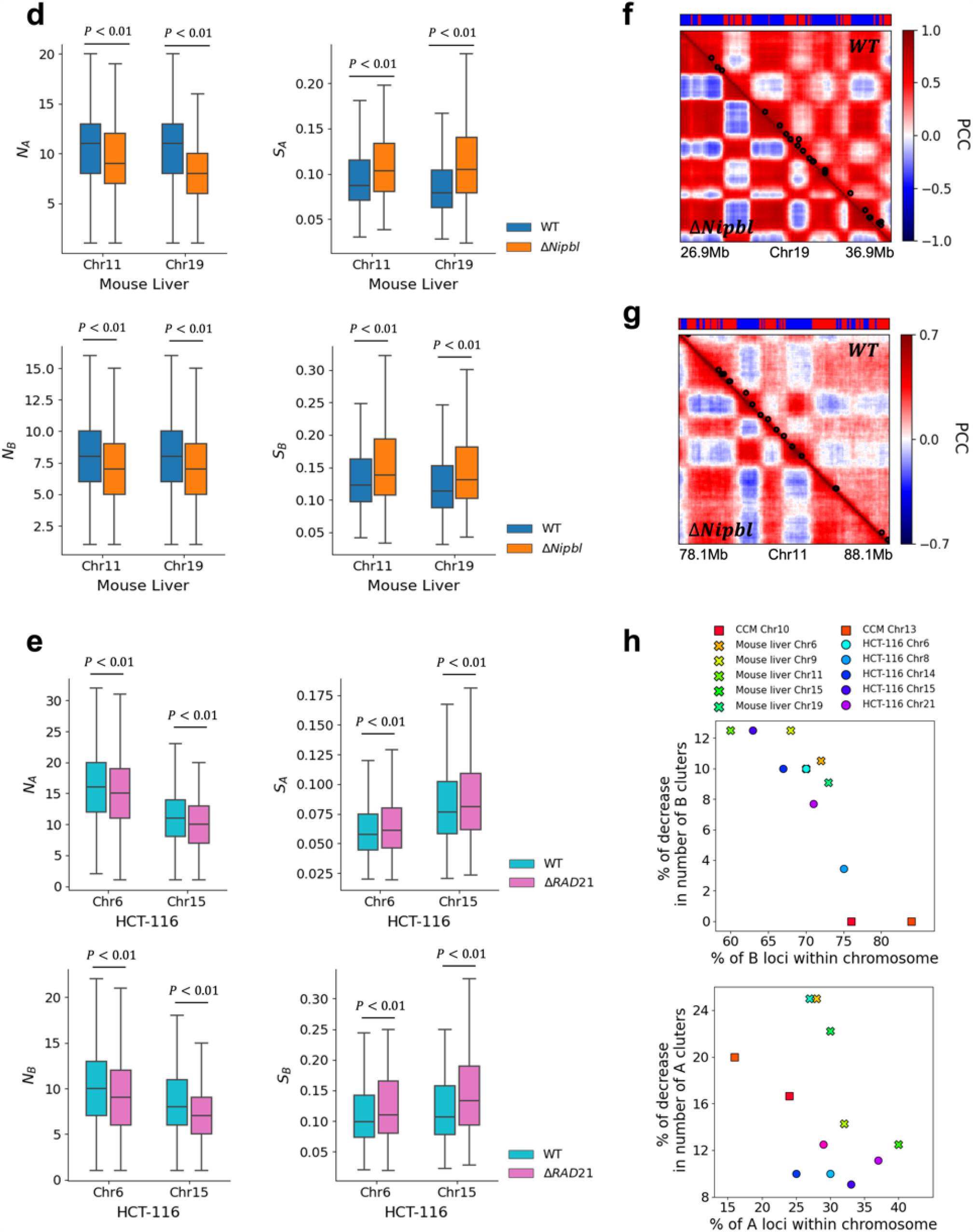
Clustering of A and B loci is stronger after loop (cohesin) loss: (a) Comparison between simulated contact maps using CCM (19-34Mb, upper panel) and Pearson correlation maps (19-29Mb, lower panel) for Chr13 (GM12878 cell line). Upper triangle (lower triangle) was calculated with (without) CTCF loops. The black circles in the upper triangle are the positions of the CTCF loop anchors detected in the Hi-C experiment [4]. The bar on top marks the epigenetic states with red (blue) representing active (repressive) loci. Upon CTCF loop loss, the plaid patterns are more prominent, and finer details of the compartment organization emerge. (b) 3D snapshots of A and B clusters identified using the DBSCAN algorithm with *P*_L_ = 1 (left panel) and *P*_L_ = 0 (right panel) computed from simulations of Chr13 with and without loops, respectively. Five A clusters (Upper panel; Red, orange, yellow, dark-green, light-green) and one B cluster (Lower panel; white) are detected in this 3D structure with *P*_L_ =1. Four A clusters and one B cluster are detected for *P*_L_ =0. The size of a locus σ _50K_ ≈ 243nm[11]. (c) Box plot of the number (left) and average size (right) of A (B) clusters determined using 10,000 individual 3D structures for *P*_L_ =1 and *P*_L_ =0 for simulated Chr10 and Chr13. The size of the A (B) cluster, *S*_A_ (*S*_B_) is defined as, (the number of A (B) loci within the cluster)/(the total number of A (B) loci within the chromosome). Boxes depict median and quartiles. The black line with caps describes the range of values in the number and size. Loop loss creates a smaller number (enhancement in compartment strength) of A-type clusters whose sizes are larger (Upper). Two-sided Mann-Whitney U test was performed for the statistical analysis. There is no change in the number and size of B clusters after loop deletion (Lower). (d)-(e) Same as (c) except the results were determined using 10,000 3D structures generated with the HIPPS method from the experimental Chr11 and Chr19 contact maps (Chr6 and Chr15 contact maps) from mouse liver for the WT and Δ *Nipbl* [34] (HCT-116 in (WT) and Δ*RAD21* cells [26]), respectively. The number of A clusters decreases by 18% and 27% after *Nipbl* loss in Chr11 and Chr19, respectively. (f)-(g) Pearson correlation matrix derived from 3D structures for Chr11 and Chr19 of mouse liver, respectively. Two loci, separated by a distance smaller than 1.75σ are in contact (σ is the mean distance between *i* and *i* + 1 loci for WT and Δ *Nipbl*, respectively). The black circles in the upper triangle are loop anchors detected in Hi-C map [34] using HiCCUPS [4]. (h) The percentage of decrease in the number of A (B) clusters after CTCF loop or cohesin loss for some chromosomes in simulations and experiments as a function of the percentage of A (B) loci within the chromosome. When the proportion of B loci is much larger than A loci, there is no change in B clusters despite loop or cohesin deletion (Upper panel).

**Appendix Fig. 5.**
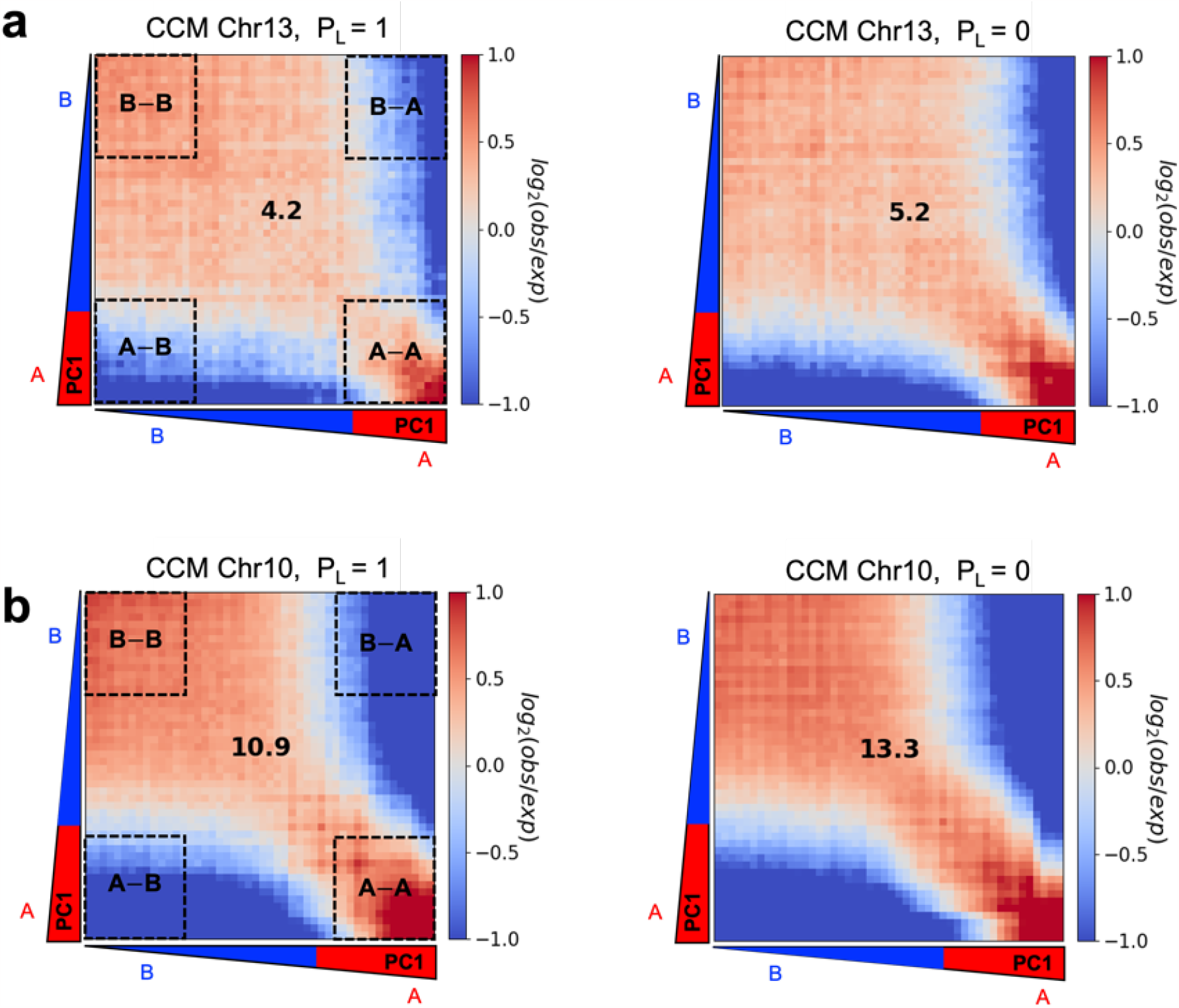
Enhancement of compartmentalization upon CTCF loop loss: Compartmentalization saddle plots are shown for (a) Chr13 and (b) Chr10 with *P*_L_ = 1 (left) and *P*_L_ = 0 (right). Observed/expected matrix bins are arranged based on PC1, obtained from the contact maps without loops. Numbers at the centre of the maps represent compartment strengths defined as the ratio of ((AA) and (BB) interactions) to ((AB) and (BA) interactions) using the mean values from the corners. The increase in the compartment score (4.2 to 5.2 for Chr13 and 10.9 to 13.3 for Chr10) shows that the compartment features are accentuated in *P*_L_ = 0 (loop deletion) compared to *P*_L_ = 1, which accords well with the conclusions in the main text that uses a different method.

**Appendix Fig. 6.**
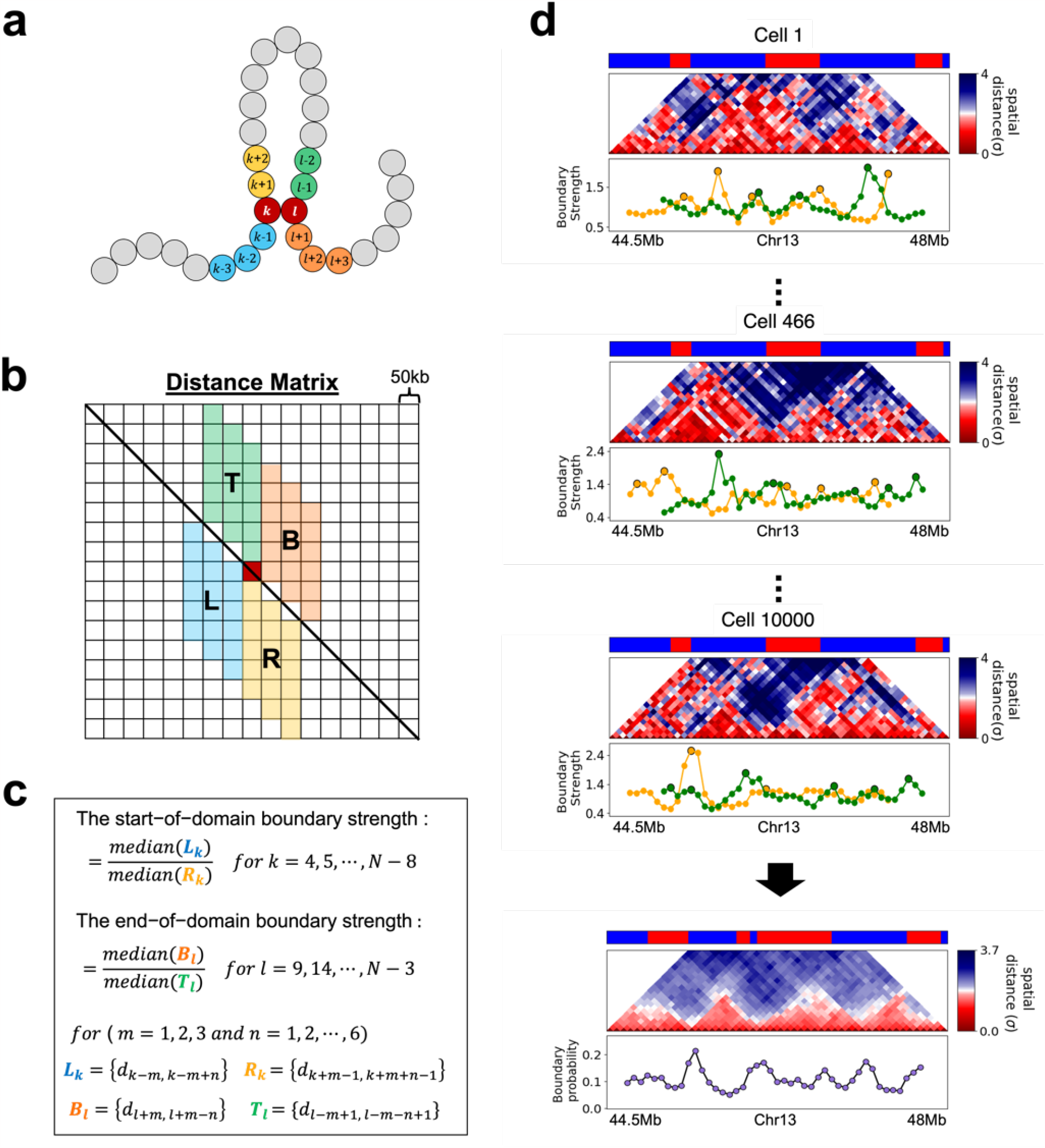
Calculation of boundary strength and boundary probability from the distance matrix at 50kb resolution: (a) A schematic describing the chromosome model. (b) Each small square of size *a* (= 50 kb) represents distance, *r*_*ij*_ between two loci *i* and *j*. The red square is used to illustrate the idea. (c) Definition of the start and end-of domain boundary strengths in the N×N distance matrix. The distance between the loci are represented as arcs in various colors. (d) The distance maps in 10,000 cells are calculated using the 3D structures using the HIPPS method[68] with Hi-C contact map from Schwarzer et al. [34] as input. Local maxima above a defined threshold at the start/end-of domain boundary strengths (yellow and green lines, respectively) are defined as domain boundaries in the WT Chr13. The start/end boundary probabilities for each locus are calculated as the proportion of cells in which the corresponding locus is a boundary location. The average of the start and end boundary probabilities cover 10,000 cells, is defined as the boundary probability for a given locus.

**Appendix Fig. 7.**
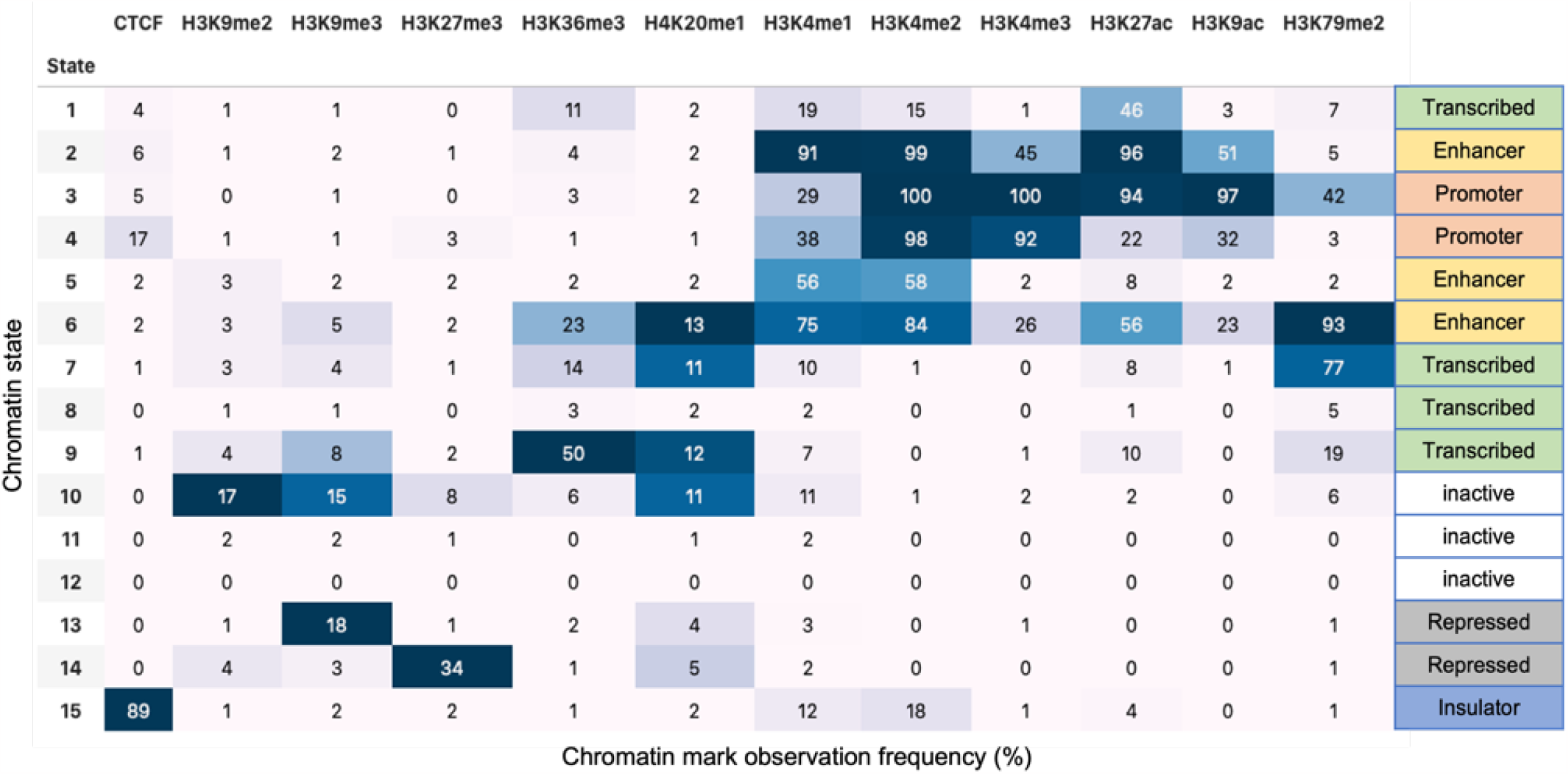
ChromHMM chromatin state annotation in HCT-116 cells

**Appendix Fig. 8.**
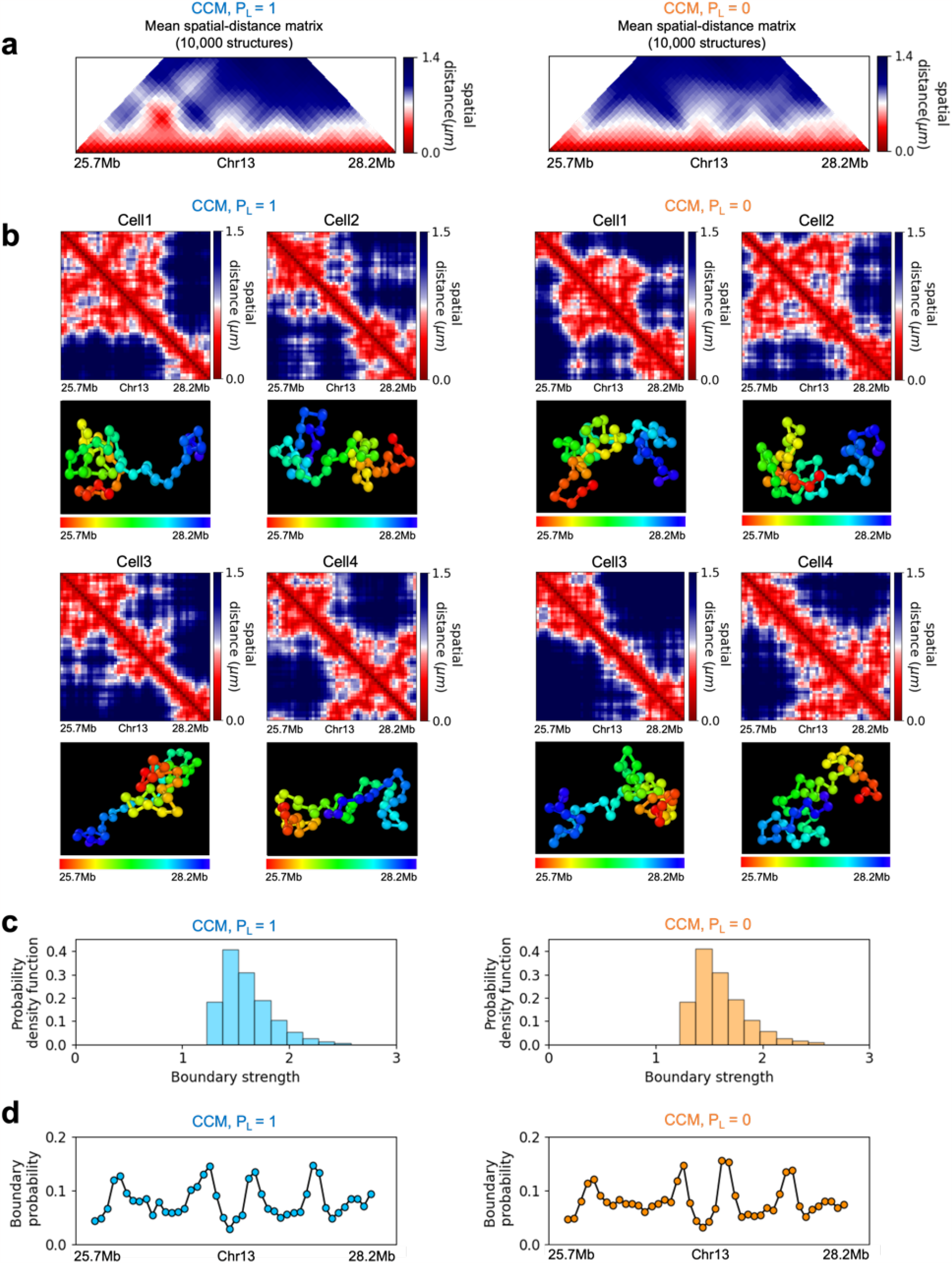
Single cell TAD-like structures exhibit in both *P*_L_ = 1 and *P*_*L*_ = 0. (a) Mean-spatial distance matrix for the genomic region (25.7-28.2Mbps) in CCM Chr13 without (left) and with (right) CTCF loops. (b) Examples of single-cell spatial-distance matrices calculated from the simulated 3D structures. TAD-like structures vary from cell to cell in both *P*_*L*_ = 1 (left) and *P*_*L*_ = 0 (right). Schematic of structures for the four cells under the two conditions are given below. (c) Distribution of the boundary strengths before (left) and after (right) CTCF loop loss, describing the steepness in the changes in the spatial distance across the boundaries. (d) The probability for each locus to be a single-cell domain boundary in cells for *P*_*L*_ = 1 (left) and *P*_*L*_ = 0(right).

**Appendix Fig. 9.**
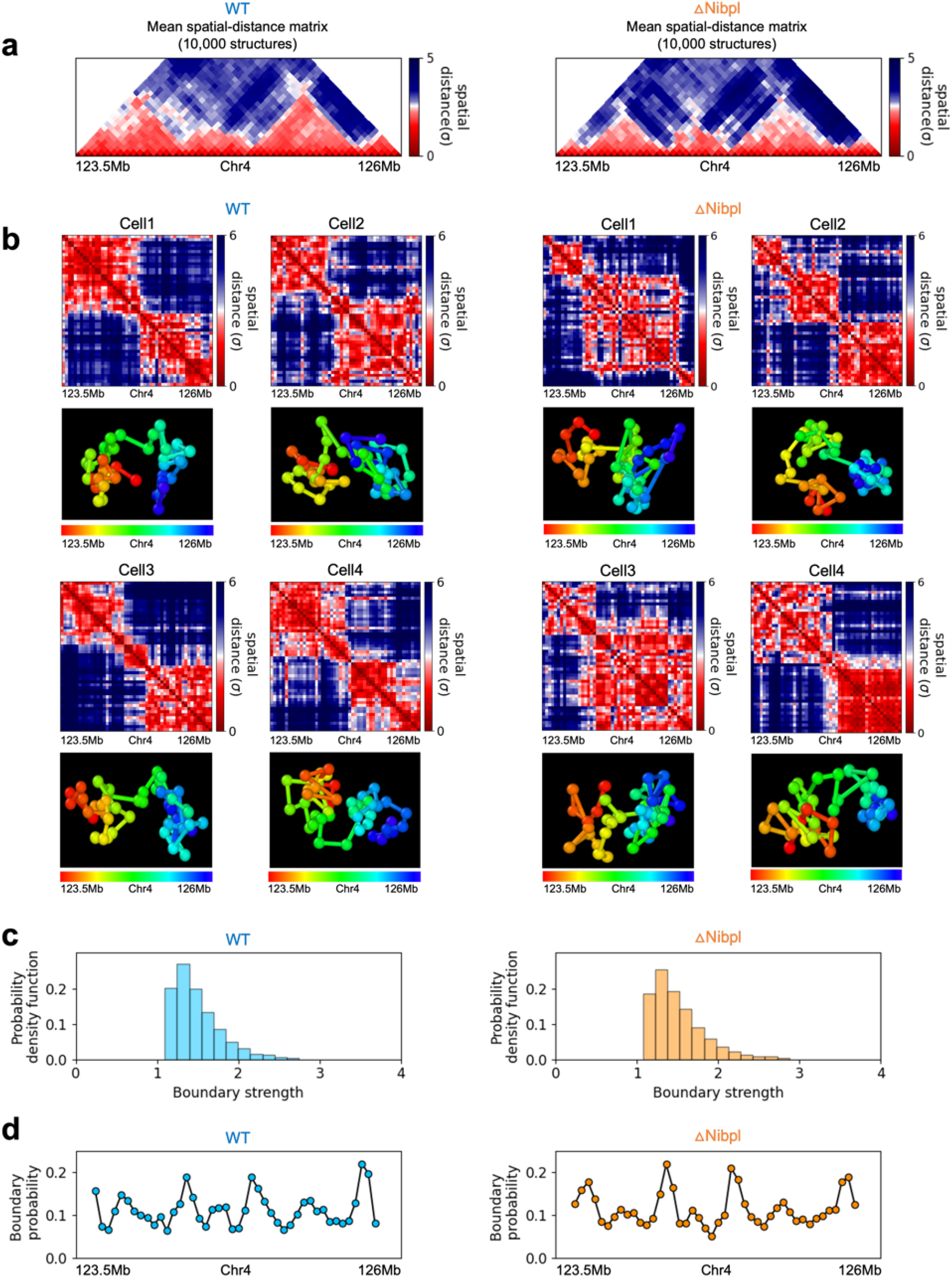
Same as Appendix Fig. 8 except the results are for the genomic region (123.5-126Mb) in Chr4 of mouse liver [34] with (left) and without (right) cohesin loading factor *Nipbl*. HIPPS generated single-cell spatial-distance matrices using Hi-C contact maps as inputs.

**Appendix Fig. 10.**
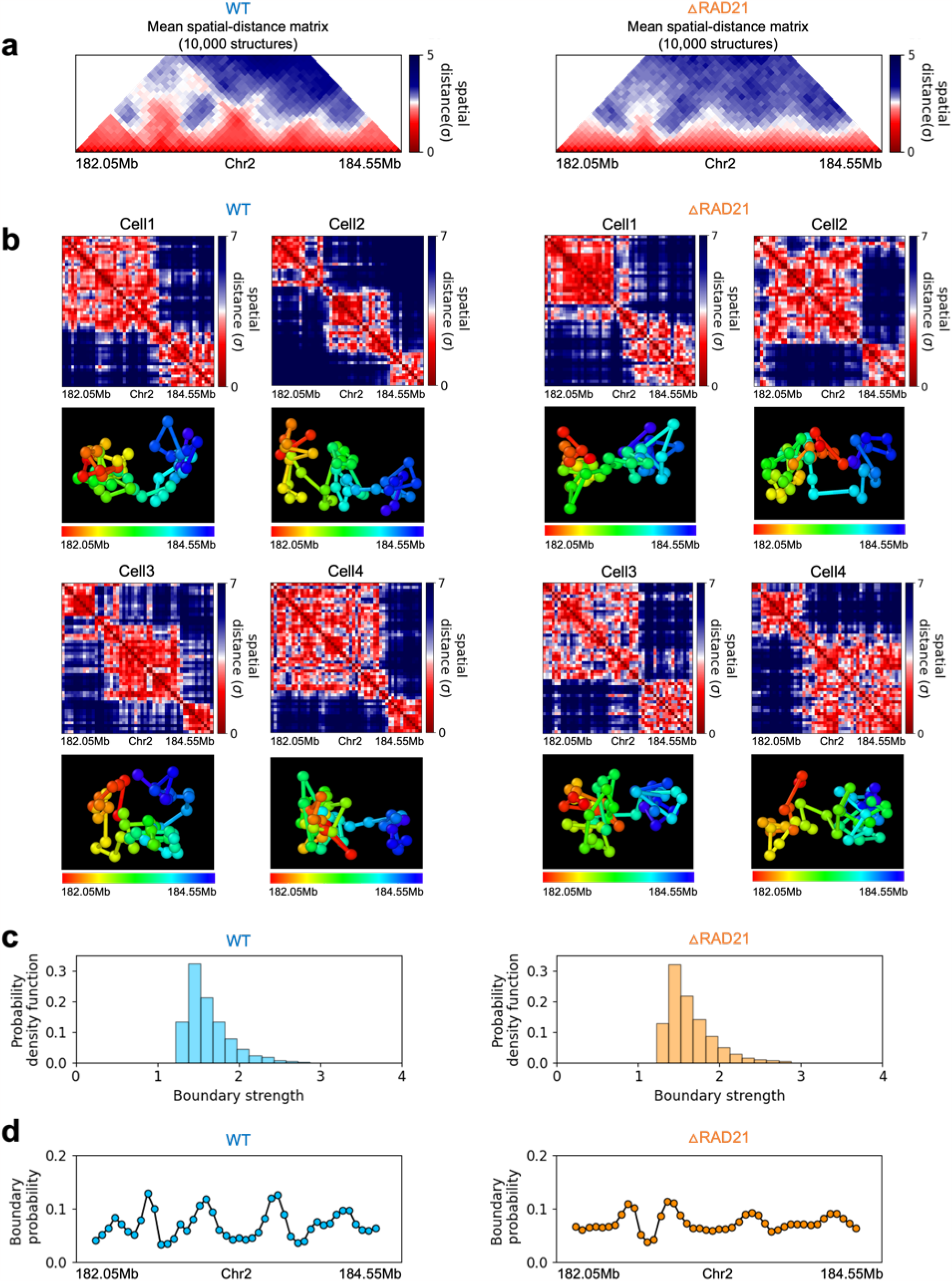
Same as Appendix Fig. 8 except the results are for the genomic region (182.05-184.55Mb) in Chr2 of HCT116 [26] with (left) and without (right) a core component of the cohesin complex, *RAD21*. Single cell 3D structures were calculated from Hi-C contact maps using HIPPS.

**Appendix Fig. 11.**
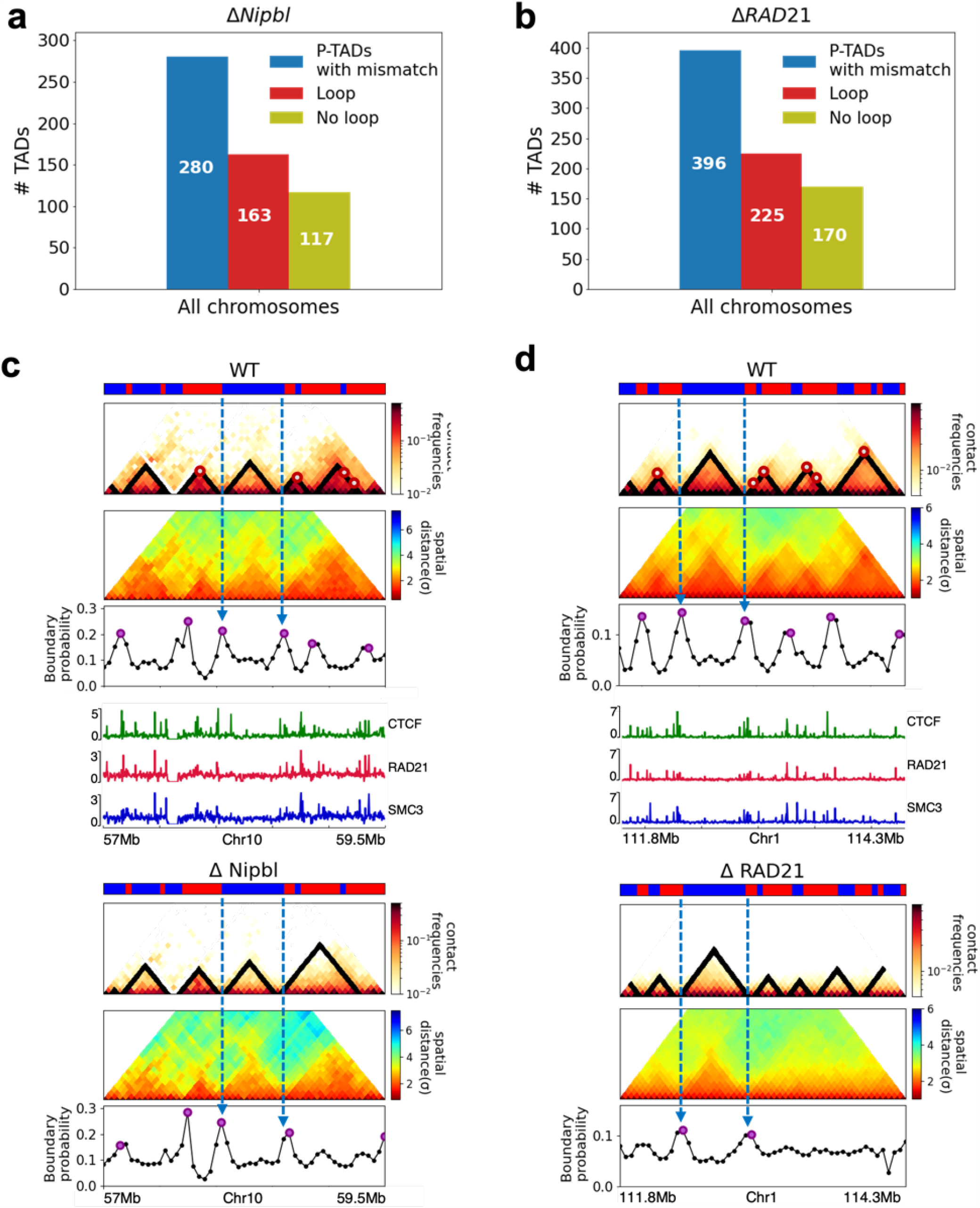
Epigenetic states contribute to the formation of domain boundaries. P-TAD does not always have corner dots at their boundaries in the WT cells. (a) The number of P-TADs (after Δ *Nipbl*) whose boundaries coincide with both epigenetic switches and corner dots (CTCF loop anchors) (red color) and only epigenetic switches (olive color) in the WT chromosomes from mouse liver. (b) Same as (a) except the results are obtained using experimental data from HCT-116 cell. (c) Chr10:57Mb-59.5Mb in mouse liver and (d) Chr1:111.8Mb-114.3Mb in HCT-116 cells, respectively. Comparison between 50kb-resolution contact maps for the 2.5Mb region with (upper) and without (lower) *Nipbl* (*RAD21*). The panels below show the mean distance maps obtained from the 3D structures. ChIP-seq tracks for CTCF, RAD21 and SMC1 in WT cells [4, 34] illustrate the correspondence between the locations of the detected loop anchors and the ChIP-seq signals. Comparison of the contact maps and boundary probabilities in (c) and (d) shows that the P-TAD boundaries (blue dotted lines) correspond well with epigenetic switch (blue line) even without corner dots in WT cells. Purple circles in the boundary probability graph represent the preferred boundaries.

**Appendix Fig. 12.**
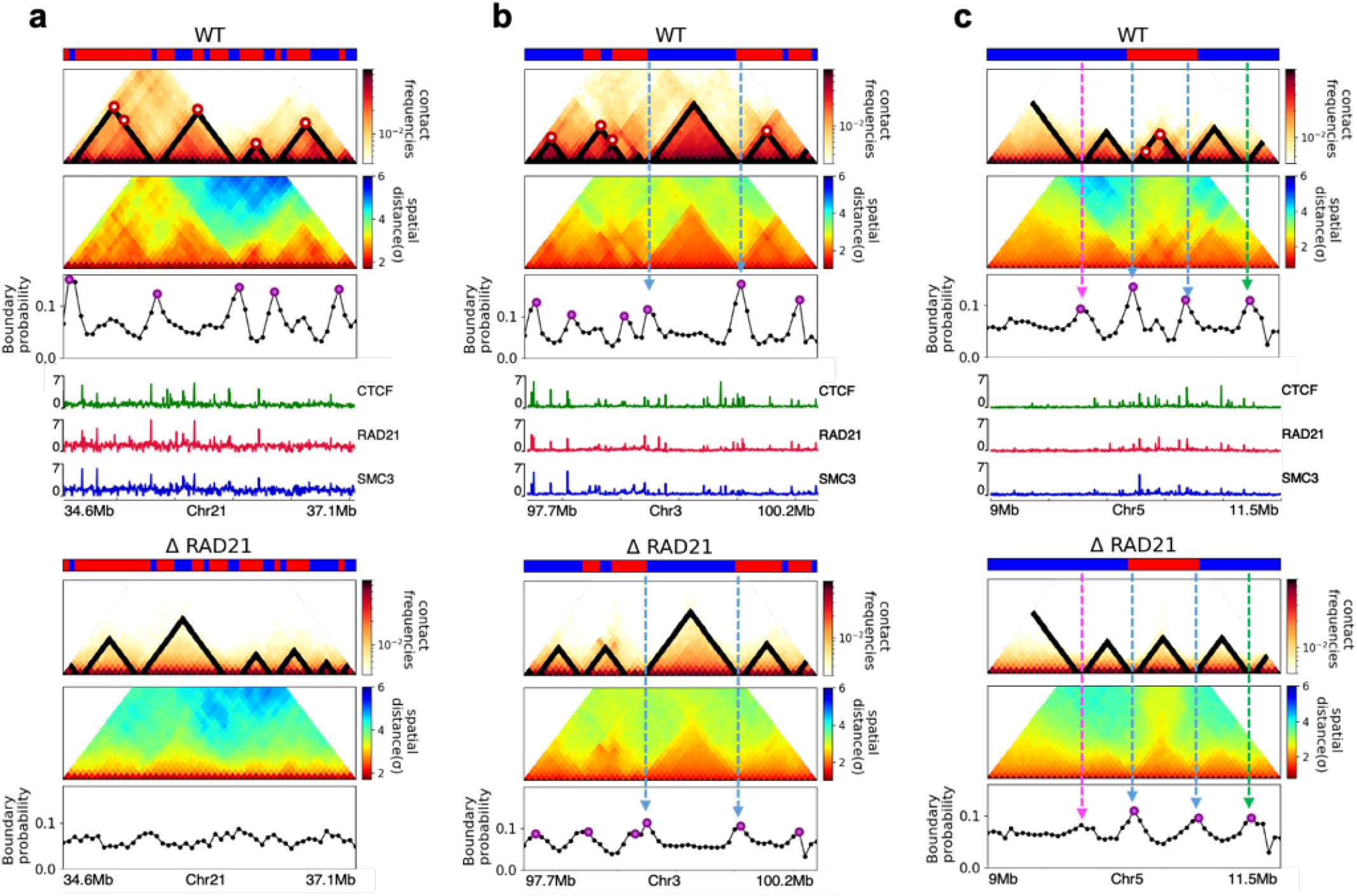
Fate of TAD structures after loss of *RAD21* in HCT-116 cells. (a) Complete Loss (Chr21:34.6Mb-37.1Mb) (b)-(c) Preserved (Chr3:97.7Mb-100.2Mb and Chr5:9Mb-11.5Mb). 50kb-resolution contact maps for the 2.5Mb genomic regions of interest with (upper) and without (lower) *RAD21* are shown in the middle panels. The dark-red circles at the boundaries of the TADs in the contact maps are loop anchors detected using HiCCUPS [55]. The mean distance maps calculated using the 3D structures with and without *RAD21* are compared in the top and bottom panels. ChIP-seq tracks for CTCF, RAD21 and SMC1 in WT cells [4] illustrate the correspondence between the locations of the detected loop anchors and the ChIP-seq signals. Bottom plots are the probability for each genomic position to be a single-cell domain boundary in the regions for cells. Purple circles in the boundary probability graph represent the preferred boundaries. Some P-TADs boundaries match epigenetic switch (blue lines). P-TADs have only high peaks in boundary probability (green line) without evidence for epigenetic switch. The magenta line shows discordance between TopDom and Boundary probability.

**Appendix Fig. 13.**
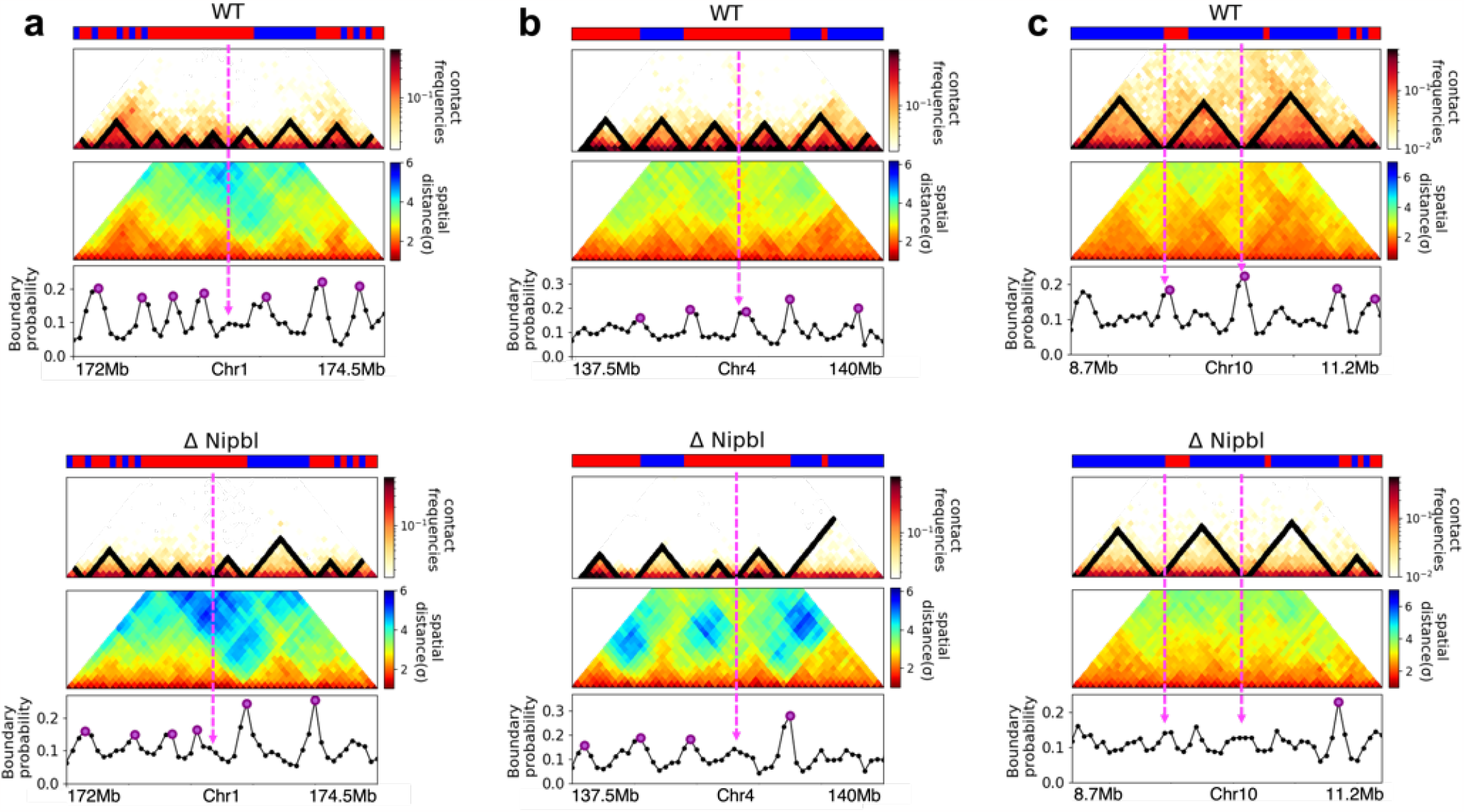
Examples of discordance between TopDom and boundary probability predictions in mouse liver [34]. In all cases, the plots show contact maps with TopDom results, mean spatial-distance matrix and boundary probability for the 2.5Mb region (a) (Chr1:172Mb-174.5Mb) (b) (Chr4:137.5Mb-140Mb) and (c) (Chr10:8.7Mb-11.2Mb) with (top) and without (bottom) *Nipbl*. Purple circles in the boundary probability indicate the prominent physical boundary in 3D structures. The magenta lines represent discordance between TopDom and Boundary probability

